# Ragnarok: a flexible and RApid GeNe Annotation (ROcKs) pipeline deployed through Nextflow

**DOI:** 10.1101/2025.10.03.680343

**Authors:** Ryan D. Kuster, Zane C. Smith, Lauren Whitt, Margaret Staton, Ben N. Mansfeld, Christopher Gottschalk

## Abstract

**Motivation:** High-quality genome assemblies and pangenomes are increasingly accessible and achievable due to advances in third-generation sequencing and assembly algorithms, but genome annotation remains a critical bottleneck. Existing gene annotation pipelines often require complex installations, multiple steps, long runtimes, and produce variable results, which impede quality and the downstream usage of the annotations.

**Results:** We developed RAGNAROK (RApid GeNe Annotation ROcKs), a modular annotation pipeline built on Nextflow and Apptainer that integrates *ab initio* prediction, transcriptome and protein evidence, repeat annotation, functional assignment, and quality assessment into a single reproducible workflow. Moreover, RAGNAROK utilizes GPU acceleration and parallelization to produce high-quality gene annotations. We benchmarked against BRAKER3 across five diverse, reference-quality plant genomes and demonstrated that RAGNAROK achieved higher sensitivity, precision, and F1 scores at the exon, transcript, and gene levels. Furthermore, we demonstrated RAGNAROK’s improvement in re-annotating a suite of five Rosaceae genomes that were previously annotated using MAKER. Overall, RAGNAROK produced more ideal mono:multi-exonic gene ratios, improved BUSCO completeness scores, and reduced missing gene content compared to other pipelines. Additionally, RAGNAROK consistently outperformed BRAKER3 in runtime, scaling efficiently from small to gigabase-scale genomes. RAGNAROK provides a flexible, rapid, scalable, and accurate solution for *de novo* and re-annotation of plant genomes. Its modular design and workflow scalability lay the foundation for future extensions to animal, fungal, and other eukaryotic genomes.

**Availability and Implementation:** RAGNAROK is available as a GitHub repository at https://github.com/ryandkuster/ragnarok.

## Introduction

With recent advancements in third-generation sequencing technologies and improved genome assembly software, the ability to assemble high-quality eukaryotic genomes has become increasingly accessible and achievable (Li and Durbin, 2024). These advances have rapidly increased genomic resources for species with large, heterozygous, and repetitive genomes, including plants (Li and Harkess, 2018; Gladman *et al*., 2023). However, genome assembly is only the first stage of generating a high-quality genomic resource. Annotation of genic and repeat-rich regions is currently the most time-intensive and computationally demanding step (Jung *et al*., 2020; Kim and Kim, 2022; Dominguez Del Angel *et al*., 2018). The requirements of annotation are amplified in the pan-genomic context, where uniformly annotating several to tens of complex genomes becomes crucial. Despite recent advances, producing accurate, reproducible, and accessible genome annotations remains a challenge.

The process of genome annotation begins within the intergenic space, which consists of low-complexity sequences that harbor transposable elements (TE), simple sequence repeats, pseudogenes, and other non-coding features. Many tools have been developed to annotate specific features within genomes, such as Generic Repeat Find, LTR_FINDER, and HelitronScanner (Xiong *et al*., 2014; Xu and Wang, 2007; Shi and Liang, 2019). Outputs of these repeat-identifying tools are often used as inputs into other software, such as LTR_retriever and RepeatModeler2 (Ou and Jiang, 2018; Flynn *et al*., 2020), to broaden their utility by identifying new or more complex related features. Over time, all-in-one pipelines or online servers that rely on such software have become available, providing greater ease-of-use (Ou *et al*., 2019; Humann *et al*., 2019; Su *et al*., 2021; Baril *et al*., 2024; The Galaxy Community, 2024). Apart from the biological importance of TE annotation, the accurate annotation of repetitive space is critical for gene annotation, as it allows masking of repeats that would otherwise contain thousands of untranslated pseudogenes, generated from TE activity in the genome. Repeat masking is typically handled by either the TE annotation pipelines themselves (*e*.*g*., EDTA) or by a separate program such as RepeatMasker (Ou *et al*., 2019; Su *et al*., 2021; Smit *et al*., 2013).

While the sensitivity and precision of TE annotators are critical to annotating a genome, the success of this step hinges on appropriate masking following TE identification. Masking typically comes in two forms, hard- or soft-masking. Hard masking replaces the nucleotides of the annotated repeats with Ns, whereas soft-masking swaps the capitalized nucleotides of the FASTA format with lowercase. Additional methods include partial hard-masking, in which hundreds to thousands of nucleotides in the left and right flanks of TEs remain unmasked in an attempt to balance between over- and under-masking. However, masking in its forms may introduce additional bias and could hinder annotating important genes or more complex features (Bayer *et al*., 2020). For example, nucleotide-binding leucine-rich repeat genes (commonly referred to as *R* genes) contain repetitive sequences intrinsically, while also commonly grouped in repetitive clusters within repeat-rich regions (Young, 2000; Hammond-Kosack and Jones, 1997; Ellis *et al*., 2000; Bayer *et al*., 2018). These arrays can be further confounded by the presence of sequence-similar pseudogenes, which can serve as regulatory elements (Balakirev and Ayala, 2003; Pink *et al*., 2011). This complexity within perceived low-complexity regions creates additional challenges to gene annotation. To overcome these challenges, *R* gene-specific annotation pipelines such as NLR-parser, NLR-Annotator, and FindPlantNLRs have been developed (Chen *et al*., 2023a; Steuernagel *et al*., 2015, 2020). Typically, these pipelines are executed on an unmasked genomic sequence to identify *R* genes. However, these tools have not yet been integrated into mainstream gene annotation pipelines, despite their ability to identify novel *R* genes missing from traditional gene annotations (Larson *et al*., 2025; Liu *et al*., 2024).

While low-complexity repetitive space remains challenging to annotate, high-complexity genic space is even more onerous. Many tools have been developed to recognize sequence features of protein-coding genes and predict gene annotations without evidence, such as Augustus, GeneMark, SNAP, and Helixer (Brůna *et al*., 2024; Stanke *et al*., 2004; Korf, 2004; Stiehler *et al*., 2021). These *ab initio* gene predictors can improve their accuracy and sensitivity for identifying features when iteratively trained (*i*.*e*., Augustus, GeneMark, and SNAP) (Stanke *et al*., 2004; Korf, 2004; Brůna *et al*., 2024). However, this process is time-consuming and requires substantial user input and operations. In comparison, some *ab initio* predictors, such as Helixer, rely on a preset inference dataset and require execution once (Stiehler *et al*., 2021). In addition to *ab initio* predictions, annotation can also leverage evidence-based inputs such as a *de novo* or reference-guided transcriptome assembly. Transcriptome assemblers, such as Trinity and StringTie2, can assemble transcriptional models from short-read RNA-seq data either *de novo* or by aligning the RNA-seq reads to the genome, respectively (Pertea *et al*., 2016; Grabherr *et al*., 2011). With the advent of third-generation sequencers, long-read RNA-seq (*i*.*e*., iso-seq) can also be employed in transcriptome assembly by programs such as StringTie2 and RNA-bloom2 (Kovaka *et al*., 2019; Nip *et al*., 2023; Shumate *et al*., 2022).

Beyond transcripts, protein sequence-based evidence annotation is also useful. Tools such as Exonerate’s protein2genome and miniprot are capable and fast protein aligners (Li, 2023; Slater and Birney, 2005). Ultimately, most published genome reports utilize a combination of *ab initio* predictions and evidence-based annotations. This combination method for gene annotation has been integrated into all-in-one annotation pipelines such as MAKER, BRAKER, and FUNANNOTATE (Campbell *et al*., 2014; Palmer and Stajich, 2020; Gabriel, Brůna, *et al*., 2024).

Although aimed at simplifying the genome annotation process, these all-in-one annotation pipelines can be challenging to install, cumbersome to execute, complex to parameterize, slow to run, and difficult to reproduce. For example, *de novo* annotations using BRAKER, where RNA-seq and protein data are available, require genome annotations generated independently by BRAKER1 (RNA-seq) and BRAKER2 (protein), followed by the transcript selection tool TSEBRA (Gabriel *et al*., 2021). However, this procedure has since been integrated into a singular pipeline in BRAKER3 (Gabriel, Brůna, *et al*., 2024). Furthermore, under- and over-annotation can be observed when using these pipelines. For example, the MAKER and BRAKER pipelines are often challenged by large, highly repetitive plant genomes and a lack of available evidence-based data (Vuruputoor *et al*., 2023). Vuruputoor et al. (2023) demonstrated these flaws by reannotating the repetitive and large *Liriodendron* genome using various combinations of RNA-seq evidence in the BRAKER and BRAKER + TSBERA pipelines. Vuruputoor et al. (2023) reported a total of 30,035 to 51,804 annotated genes, compared to the published annotation of 35,261, when using different RNA-seq inputs and filtering methods (Vuruputoor *et al*., 2023; Chen *et al*., 2019), demonstrating the high variability in annotation.

To overcome the challenges associated with annotation, we developed a Nextflow pipeline (Langer *et al*., 2025) that utilizes Apptainer (Kurtzer *et al*., 2017) to combine a suite of fast and accurate, *ab initio* and evidence-based gene annotators into a novel, all-in-one annotation pipeline (Langer *et al*., 2025). This pipeline, named RApid GeNe Annotation (ROcKs) or RAGNAROK (https://github.com/ryandkuster/ragnarok), aims to be a rapid, flexible, accurate, and reproducible option for *de novo* and re-annotation of plant genomes. Moreover, RAGNAROK’s implementation also allows for the optional incorporation of repeat annotation using EDTA before gene annotation. Furthermore, the pipeline performs integrated gene model filtering, functional annotation, and quality assessment. We benchmarked our pipeline, RAGNAROK, against other annotation pipelines (MAKER and BRAKER) on both model and non-model genomes to demonstrate its higher accuracy, precision, and overall annotation quality. By deploying our pipeline through the increasingly popular Nextflow workflow management system (Langer *et al*., 2025), we eliminate the need for users to install all dependencies, thereby creating a reproducible and scalable workflow.

## Methods

### Pipeline development and implementation

RAGNAROK annotation is split into several parallel processes for annotation: 1) *ab initio* gene prediction, 2) evidence-based transcriptome assembly, 3) protein database mapping, and 4) previous annotation liftover (Fig. 1). For *ab initio* gene prediction, RAGNAROK leverages the graphical processing unit (GPU) accelerated Helixer gene predictor (Stiehler *et al*., 2021), which maintains support for the CPU implementation of Helixer at the expense of additional processing time. Here, the masked genome file and the Helixer scoring file are used as inputs into Helixer with default plant parameters (Stiehler *et al*., 2021; Holst *et al*., 2023).

**Figure 1.**
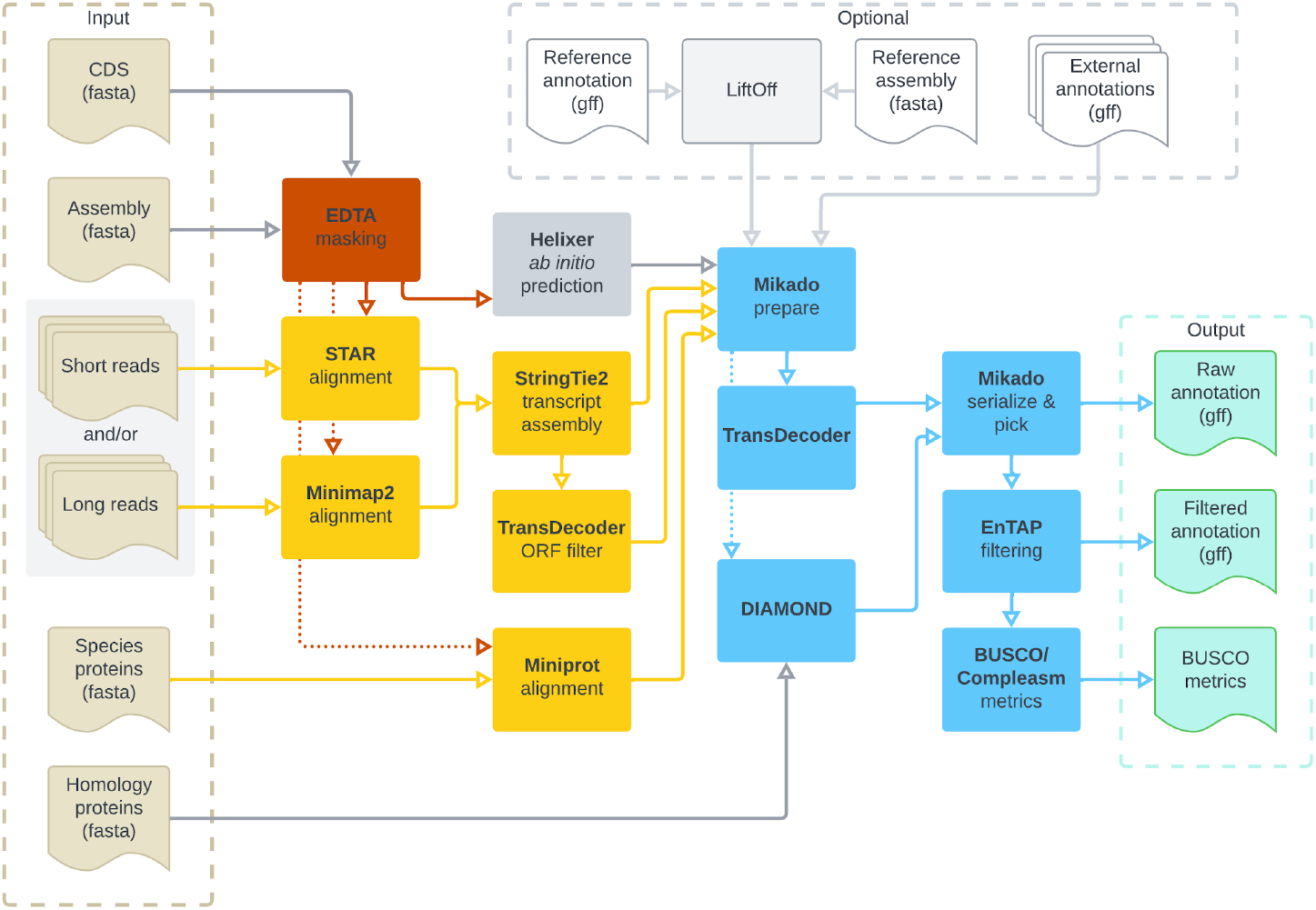
Flowchart of critical RAGNAROK pipeline stages. Evidence-based steps are represented in yellow, and all steps post-Mikado are represented in blue.

For evidence-based transcriptome assembly, we utilize the StringTie2 pipeline with short- and/or long-reads (Pertea *et al*., 2016; Kovaka *et al*., 2019; Shumate *et al*., 2022). With short-read data, reads are first assessed for quality using FASTQC and MULTIQC, and processed for adapters and trimmed by FASTP (Chen *et al*., 2018; Andrews, 2010; Ewels *et al*., 2016). A second assessment of the resulting trimmed reads is conducted with FASTQC and MULTIQC. The masked genome file is then indexed using the STAR aligner, and trimmed reads are mapped onto the genome using the following parameters --outSAMtype BAM Unsorted --outSAMstrandField intronMotif --alignIntronMax 10000 (Dobin *et al*., 2013). With long-read data, the masked genome is indexed using minimap2, and reads are aligned with the following parameters -ax splice:hq -uf enabled (Li, 2018). All aligned reads are then sorted using SAMtools and used as input into StringTie2, with the hybrid mode enabled if long reads were used, to perform transcript assembly (Pertea *et al*., 2016; Kovaka *et al*., 2019; Shumate *et al*., 2022; Danecek *et al*., 2021; Pertea *et al*., 2015). The resulting GTF annotation file is subsequently converted to GFF3 format using gffread (Pertea and Pertea, 2020). We additionally use the StringTie2 GTF as input into TransDecoder (https://github.com/TransDecoder/). TransDecoder aims to extract the longest open-reading frame transcripts from the StringTie2 output. Here, we first execute gtf_genome_to_cdna_fasta.pl using the StringTie2 GTF and masked genome file. We then execute *TransDecoder*.*LongOrfs* and *TransDecoder*.*Predict* using the output from gtf_genome_to_cdna_fasta.pl. The StringTie2 GTF is then used again as input into *gtf_to_alignment_gff3*.*pl*. The last step of the TransDecoder steps was *cdna_alignment_orf_to_genome_orf*.*pl* using the outputs from all three prior steps as suggested by TransDecoder’s documentation (https://github.com/TransDecoder/).

Our second evidence-based annotation comes from alignments of a reference protein data set using miniprot (Li, 2023). First, an index is created from the masked genome file. Next, the reference protein fasta is used as input into miniprot with the flag for –gff evoked for output (Li, 2023). We also added an additional, optional step for evidence-based annotation via LiftOff (1.6.3) to lift over existing reference annotations from a similar species by selecting optimal exon alignments (Shumate and Salzberg, 2021). LiftOff is also particularly useful for transferring pre-existing gene annotations from previous annotation versions. This step was executed using the default parameters and gene copy search, considering only alignments with at least 95% identity. Additionally, RAGNAROK flexibly supports the inclusion of user-generated GFFs from other tools, *e*.*g*., cufflinks (Trapnell *et al*., 2012) or prothint (Brůna *et al*., 2020), or other annotation pipelines. However, the user must include their own weights for Mikado transcript selection.

Following gene predictions and evidence-based annotations, we use the Mikado transcript selector to reduce redundancy and select the best representative transcript models (Venturini *et al*., 2018). Although written for polishing of reference annotations, Mikado serves usefully in selecting a consensus annotation from *de novo* and *ab initio* sources. The first step in transcript selection by Mikado was the design of the input list file. This tab-delimited file includes all the resulting GTF/GFFs from all previous annotation steps as described by Mikado’s documentation. These inputs included the StringTie2 GTF, miniprot GFF, Helixer GFF, TransDecoder GFF, and optionally the LiftOff GFF. The miniprot, StringTie2, and LiftOff datasets were flagged as reference annotations. Our scoring for all inputs was scaled by our confidence in the annotation tools (StringTie2=10, miniprot=10, Helixer=1, TransDecoder=-0.5). With the list file created, we then executed Mikado configure with the list file as the –list input, the masked genome file as the –reference input, the Mikado prepared plants.yaml as the –scoring input, the UniProt Swiss-Prot protein sequences (The UniProt Consortium, 2025) as the –bt input, and the Mikado provided configuration.yaml file as the final input. We then executed mikado prepare --json-conf using the mikado prepared configuration.yaml (Venturini *et al*., 2018). Once completed, we then prepared a DIAMOND database using the UniProt Swiss-Prot proteins (Buchfink *et al*., 2021). DIAMOND was then executed on the Mikado prepared transcript fasta file with the following flags: blastx -k 5 -f 6 qseqid sseqid pident length mismatch gapopen qstart qend sstart send evalue bitscore ppos btop (Buchfink *et al*., 2021). Next, we used TransDecoder.LongOrfs on the Mikado prepared transcript fasta file. We were then able to continue with the Mikado pipeline. First, Mikado serialise was executed using the Mikado prepared configuration.yaml as the --json-conf input, the Mikado prepared fasta as --transcripts input, the DIAMOND output at the --xml input, and the TransDecoder.LongOrfs output was prepared on the Mikado transcript fasta as the --orfs input, and finally, the UniProt Swiss-Prot proteins were used as the --blast_targets input (Venturini *et al*., 2018). The final step was to execute the Mikado pick with the masked genome fasta as the --fasta input, Mikado prepared the configuration.yaml as the --configuration input, and the --subloci-out and --loci-out outputs were set. The resulting loci-out file was treated as a final, unfiltered gene annotation file (Venturini *et al*., 2018).

To further process and increase the evidence that a gene is annotated correctly, RAGNAROK carries out functional annotation. This functional annotation is conducted using EnTAP (2.2.0) (Hart *et al*., 2020). Briefly, EnTAP conducts functional annotation by performing similarity searches of the protein to several databases using DIAMOND (Buchfink *et al*., 2021). We specifically use annotations from Swiss-Prot and NCBI RefSeq Plant databases as references. Gene Ontology (GO) terms are additionally assigned using EggNOG-mapper (v.2) and the EggNOG database (v.6) (Cantalapiedra *et al*., 2021; Hernández-Plaza *et al*., 2022). Following functional annotation, we created a custom script to further filter the annotations from Mikado2 to remove pseudogenes and monoexonic features that lack transcribed proteins. This custom filtering script relies on AGAT (v1.4.1) subset functionality (Dainet, 2019).

To assess the quality of the resulting RAGNAROK annotation, we integrated several assessment tools. First, to determine the accuracy of annotating universal single-copy orthologs and utilize the benchmarking program BUSCO (5.8.2) (Simão *et al*., 2015; Manni *et al*., 2021). BUSCO is implemented to utilize the latest OrthoDB v12 databases (Tegenfeldt *et al*., 2025). Complementary, we also implemented Compleasm, which is functionally similar to BUSCO with a reportedly higher sensitivity (Huang and Li, 2023).

### Benchmarking

To evaluate the performance of RAGNAROK for *de novo* annotation, we set out to compare against BRAKER3 on several plant genomes with either heavily curated reference gene annotations or that have undergone extensive annotation parameter testing (Gabriel, Brůna, *et al*., 2024). These genomes include *Arabidopsis thaliana* TAIR10, *Prunus persica* ‘Lovell 2D’ v2, *Malus domestica* GDDH13, *Zea mays* B73, and *Liriodendron chinense* (Chen *et al*., 2019; Lamesch *et al*., 2012; Wu *et al*., 2023; Verde *et al*., 2013, 2017; Daccord *et al*., 2017; Cheng *et al*., 2017; Hufford *et al*., 2021). To directly compare against BRAKER3, we utilized the same TE annotation and evidence datasets (*e*.*g*., RNA-seq and protein databases). This was accomplished by using the EDTA mod.MAKER.masked (FASTA) output generated within RAGNAROK as masked genome input into BRAKER3, thus ensuring both pipelines annotated the same masked sequence. All corresponding commands for RAGNAROK and BRAKER3 for these five genomes can be found in Supplemental File 1. Precision and sensitivity statistics of the two resulting *de novo* annotations per species were obtained using Mikado2’s stats function and BUSCO using OrthoDB v10 and v12 (Venturini *et al*., 2018; Simão *et al*., 2015; Manni *et al*., 2021; Tegenfeldt *et al*., 2025). Comparisons between the two *de novo* annotations and the reference annotation of each species were obtained using Mikado2 compare feature (Venturini *et al*., 2018). Genomes, reference annotations, evidence datasets, and versions for each of the five subject species can be found in Supplement Table 1 (Chen *et al*., 2019; Lamesch *et al*., 2012; Verde *et al*., 2013, 2017; Daccord *et al*., 2017; Hufford *et al*., 2021).

**Table 1.** Annotation statistics between RAGANAROK, BRAKER3, and reference annotation on five reference quality genomes.

| Species | Annotation | Number of Genes | Mono-exonic Genes | Mono: Multi-exonic | Missed Exons | Matched Intron Chains | Missed Genes | Novel Genes | BUSCO Embryophyta ODB10 | BUSCO ODB12<sup>1</sup> |
| --- | --- | --- | --- | --- | --- | --- | --- | --- | --- | --- |
| *Arabidopsis thaliana* | Araport11 (reference) | 27,655 | 5,382 | 0.242 | - | - | - | - | C:99.6%[S:48.9%,D:50.7%], F:0.1%,M:0.3%,n:1614 | C:98.9%[S:48.7%,D:50.2%], F:0.4%,M:0.6%,n:2026 |
| *Arabidopsis thaliana* | BRAKER3 | 27,596 | 7,959 | 0.405 | 41,113 | 17,728 | 2,551 | 2,156 | C:98.8%[S:84.7%,D:14.1%], F:0.2%,M:0.9%,n:1614 | C:98.4%[S:83.4%,D:15.0%], F:0.5%,M:1.1%,n:2026 |
| *Arabidopsis thaliana* | RAGNAROK | 26,900 | 4,205 | 0.185 | 10,399 | 19,863 | 1,853 | 1,296 | C:98.8%[S:92.6%,D:6.2%], F:0.3%,M:0.9%,n:1614 | C:97.9%[S:89.9%,D:8.0%], F:0.6%,M:1.5%,n:2026 |
| *Prunus persica* | Prunus_persica_v2.0.a1 (reference) | 26,873 | 5,130 | 0.236 | - | - | - | - | C:99.3%[S:97.1%,D:2.2%], F:0.2%,M:0.5%,n:1614 | C:99.2%[S:99.1%,D:0.1%], F:0.2%,M:0.6%,n:10071 |
| *Prunus persica* | BRAKER3 | 30,425 | 11,531 | 0.610 | 42,250 | 14,606 | 3,435 | 6,295 | C:98.3%[S:81.7%,D:16.5%], F:0.5%,M:1.2%,n:1614 | C:98.6%[S:84.6%,D:14.1%], F:0.1%,M:1.3%,n:10071 |
| *Prunus persica* | RAGNAROK | 26,212 | 3,805 | 0.170 | 31,296 | 16,859 | 2,908 | 2,188 | C:99.1%[S:93.1%,D:6.0%], F:0.3%,M:0.6%,n:1614 | C:98.8%[S:94.1%,D:4.7%], F:0.2%,M:1.1%,n:10071 |
| *Malus x domestica* | gene_models_20170612 (reference) | 46,558 | 7,356 | 0.188 | - | - | - | - | C:96.4%[S:64.8%,D:31.6%], F:1.7%,M:1.9%,n:1614 | C:94.7%[S:36.6%,D:58.1%], F:0.3%,M:5.0%,n:10071 |
| *Malus x domestica* | BRAKER3 | 43,203 | 13,891 | 0.474 | 62,552 | 18,213 | 9,464 | 6,235 | C:97.1%[S:53.3%,D:43.9%], F:0.8%,M:2.0%,n:1614 | C:95.7%[S:28.1%,D:67.6%], F:0.1%,M:4.2%,n:10071 |
| *Malus x domestica* | RAGNAROK | 46,974 | 7,858 | 0.201 | 40,713 | 21,490 | 6,323 | 7,969 | C:98.2%[S:60.2%,D:38.0%], F:0.6%,M:1.2%,n:1614 | C:96.1%[S:32.0%,D:64.2%], F:0.2%,M:3.7%,n:10071 |
| *Liriodendron chinense* | Lich.1_0 (reference) | 35,269 | 468 | 0.013 | - | - | - | - | C:76.1%[S:69.9%,D:6.2%], F:15.2%,M:8.7%,n:1614 | C:73.8%[S:67.9%,D:6.0%], F:19.5%,M:6.6%,n:2026 |
| *Liriodendron chinense* | BRAKER3 (short-reads) | 42,083 | 19,604 | 0.872 | 111,948 | 6,398 | 12,819 | 18,248 | C:81.1%[S:60.5%,D:20.6%], F:5.0%,M:13.9%,n:1614 | C:81.3%[S:61.9%,D:19.3%], F:6.8%,M:11.9%,n:2026 |
| *Liriodendron chinense* | RAGNAROK (short-reads) | 35,223 | 3,786 | 0.120 | 96,810 | 11,343 | 10,910 | 9,379 | C:81.6%[S:72.2%,D:9.4%], F:12.4%,M:6.0%,n:1614 | C:78.7%[S:69.2%,D:9.4%], F:16.3%,M:5.0%,n:2026 |
| *Liriodendron chinense* | RAGNAROK (long-reads) | 34,500 | 3,327 | 0.107 | 95,345 | 10,612 | 11,011 | 9,250 | C:84.3%[S:76.1%,D:8.2%], F:9.5%,M:6.2%,n:1614 | C:82.4%[S:74.1%,D:8.3%], F:12.4%,M:5.2%,n:2026 |
| *Zea mays* | MaizeGDB Zm-B73 NAM-5.0 (reference) | 39,756 | 8,866 | 0.287 | - | - | - | - | C:97.0%[S:30.9%,D:66.1%], F:0.9%,M:2.1%,n:1614 | C:95.3%[S:30.3%,D:65.1%], F:2.6%,M:2.1%,n:2026 |
| *Zea mays* | BRAKER3 | 49,367 | 19,718 | 0.665 | 77,113 | 19,801 | 7209 | 15857 | C:95.6%[S:64.4%,D:31.2%], F:1.7%,M:2.7%,n:1614 | C:94.3%[S:62.9%,D:31.4%], F:3.2%,M:2.5%,n:2026 |
| *Zea mays* | RAGNAROK | 42,313 | 5,760 | 0.158 | 64,421 | 20,888 | 7433 | 9553 | C:95.5%[S:81.5%,D:14.1%], F:1.8%,M:2.7%,n:1614 | C:93.7%[S:75.1%,D:18.6%], F:3.2%,M:3.1%,n:2026 |

<sup>1</sup>For ODB12, *A. thaliana*, *L. chinense*, and *Z. mays* are reporting the embryophyta lineage scores, and *M. domestica* and *P. persica* are reporting Rosaceae.

To compare run times between RAGNAROK and BRAKER3, all five reference quality genomes were annotated in a paired fashion on the same computing hardware. For *A. thaliana, P. persica, M. domestica*, and *L. chinense*, their RAGNAROK and BRAKER3 annotations were generated on a local server. For RAGNAROK, the nextflow script was executed using the profile local and eight (SFile 1). This profile allowed for the greatest parallelization of jobs on a server outfitted with dual CPUs capable of 96 threads, 200 GB RAM, and twin NVIDIA RTX 4500 GPUs (Santa Clara, CA). For BRAKER3, all four datasets run on local hardware were executed with the threads parameter set to the maximum allowed of 48 (SFile 1). *Z. mays* was executed on a high-performance computing cluster due to the computational demands for a large, complex genome. Here, RAGNAROK was executed using the profile slurm and default settings (SFile 1). For BRAKER3, it was again set to a maximum thread parameter of 48.

We also evaluated RAGANROK for re-annotation of several non-model species that were previously annotated using the MAKER pipeline (Campbell *et al*., 2014). These genomes included *Malus angustifolia, Malus fusca, Prunus persica* ‘Lovell 2D’ v3 and ‘Lovell 5D’, and *Pyrus communis* ‘d’Anjou’ (Gottschalk *et al*., 2025; Mansfeld, Yocca, *et al*., 2023; Mansfeld, Ou, *et al*., 2023; Yocca *et al*., 2024). Similar to the *de novo* annotation, the same masked genome file and evidence datasets were used as input. Statistics for the RAGNAROK and MAKER annotations were obtained using Mikado2 stats function and BUSCO using OrthoDB v10 and v12 (Venturini *et al*., 2018; Simão *et al*., 2015; Manni *et al*., 2021; Tegenfeldt *et al*., 2025). Genomes, reference annotations, evidence datasets, and versions for each of the five subject species can also be found in Supplement Table 1. Statistical analysis was conducted using R (v.4.5.1) (R Core Team, 2024). Figure images were generated using ggplot2 (Wickham, 2016).

## Results

### Comparing de novo annotation of RAGNAROK vs BRAKER3

To evaluate the performance of RAGNAROK, we compared it directly to BRAKER3, the most widely used eukaryotic annotation pipeline (Gabriel, Brůna, *et al*., 2024). Here, we selected five reference-quality or thoroughly annotated plant genomes - *Arabidopsis thaliana, Prunus persica, Malus domestica, Zea mays, and Liriodendron chinense* to serve as independent cases. Each genome was treated as being *de novo* annotated using publicly available data that was either presented in the original publication of the genome or in previous evaluations of BRAKER3 (STable 1). To compare the quality of the annotations, the separate annotations generated by RAGNAROK and BRAKER3 were compared to the reference annotations (STable 1).

Overall, we observed higher sensitivity and precision with RAGNAROK compared to BRAKER3 at the base, exon, transcript, and gene level when compared to the reference annotation of the genome (Fig. 2; STable 2). Significantly higher sensitivity was observed for RAGNAROK at the exon, transcript, and gene levels. This trend continued for precision, with RAGNAROK being significantly higher in precision at the transcript and gene level. As a result, the F1 values, which are dependent on sensitivity and precision, were also found to be significantly higher for RAGNAROK annotations at the transcript and gene level (Fig.3; STable 2).

**Table 2.** Re-annotation statistics between MAKER and RAGNAROK.

| Species | Genome Version | Haplotype | Annotation | Number of Genes | Mono-exonic Genes | Mono: Multi-exonic | BUSCO Embryophyta obd10 | BUSCO Rosaceae obd12 |
| --- | --- | --- | --- | --- | --- | --- | --- | --- |
| *Malus angustifolia* | 1.0 | Hap1 | MAKER | 43,949 | 3,255 | 0.080 | C:95.7%[S:62.1%,D:33.6%],<br>F:1.4%,M:3.0%,n:1614 | C:89.6%[S:33.1%,D:56.5%],<br>F:0.3%,M:10.1%,n:10071 |
| *Malus angustifolia* | 1.0 | Hap2 | MAKER | 47,334 | 2,677 | 0.060 | C:95.8%[S:63.9%,D:31.8%],<br>F:1.4%,M:2.9%,n:1614 | C:89.8%[S:33.3%,D:56.5%],<br>F:0.2%,M:10.0%,n:10071 |
| *Malus angustifolia* | 1.0 | Hap1 | RAGNAROK | 48,538 | 10,117 | 0.263 | C:99.0%[S:61.3%,D:37.7%],<br>F:0.2%,M:0.7%,n:1614 | C:96.6%[S:31.7%,D:64.9%],<br>F:0.1%,M:3.3%,n:10071 |
| *Malus angustifolia* | 1.0 | Hap2 | RAGNAROK | 48,518 | 10,024 | 0.260 | C:98.8%[S:62.3%,D:36.5%],<br>F:0.4%,M:0.9%,n:1614 | C:96.6%[S:32.0%,D:64.6%],<br>F:0.1%,M:3.3%,n:10071 |
| *Malus fusca* | 1.0 | Hap1 | MAKER | 46,300 | 4,452 | 0.106 | C:96.6%[S:61.8%,D:34.8%],<br>F:1.2%,M:2.2%,n:1614 | C:93.0%[S:33.5%,D:59.5%],<br>F:0.5%,M:6.5%,n:10071 |
| *Malus fusca* | 1.0 | Hap2 | MAKER | 45,509 | 4,572 | 0.112 | C:96.3%[S:61.9%,D:34.4%],<br>F:1.2%,M:2.5%,n:1614 | C:92.9%[S:33.7%,D:59.1%],<br>F:0.4%,M:6.7%,n:10071 |
| *Malus fusca* | 1.0 | Hap1 | RAGNAROK | 42,016 | 6,272 | 0.175 | C:98.3%[S:58.8%,D:39.5%],<br>F:0.7%,M:1.0%,n:1614 | C:96.2%[S:30.9%,D:65.4%],<br>F:0.2%,M:3.5%,n:10071 |
| *Malus fusca* | 1.0 | Hap2 | RAGNAROK | 42,034 | 6,268 | 0.175 | C:98.5%[S:58.6%,D:40.0%],<br>F:0.6%,M:0.9%,n:1614 | C:96.3%[S:31.0%,D:65.3%],<br>F:0.2%,M:3.5%,n:10071 |
| *Prunus persica*<br>'Lovell 2D' | 3.0 | - | MAKER | 26,834 | 4,081 | 0.179 | C:97.0%[S:94.8%,D:2.2%],<br>F:1.8%,M:1.2%,n:1614 | C:95.3%[S:95.1%,D:0.2%],<br>F:1.0%,M:3.6%,n:10071 |
| *Prunus persica*<br>'Lovell 2D' | 3.0 | - | RAGNAROK | 31,847 | 5,371 | 0.203 | C:99.3%[S:93.3%,D:6.0%],<br>F:0.2%,M:0.5%,n:1614 | C:99.0%[S:94.0%,D:0.5%],<br>F:0.1%,M:0.9%,n:10071 |
| *Prunus persica*<br>'Lovell 5D' | 1.0 | - | MAKER | 26,764 | 3,184 | 0.135 | C:98.4%[S:92.6%,D:5.8%],<br>F:0.9%,M:0.7%,n:1614 | C:96.3%[S:92.2%,D:4.0%],<br>F:0.9%,M:2.8%,n:10071 |
| *Prunus persica*<br>'Lovell 5D' | 1.0 | - | RAGNAROK | 29,087 | 4,367 | 0.177 | C:99.2%[S:87.1%,D:12.1%],<br>F:0.3%,M:0.5%,n:1614 | C:98.9%[S:87.7%,D:11.2%],<br>F:0.2%,M:1.0%,n:10071 |
| *Pyrus communis* | 2.3.a1 | Hap1 | MAKER | 44,558 | 4,110 | 0.102 | C:98.1%[S:61.6%,D:36.5%],<br>F:0.6%,M:1.2%,n:1614 | C:94.0%[S:33.2%,D:60.8%],<br>F:0.3%,M:5.7%,n:10071 |
| *Pyrus communis* | 2.3.a1 | Hap2 | MAKER | 44,349 | 4,143 | 0.103 | C:98.4%[S:62.0%,D:36.4%],<br>F:0.5%,M:1.1%,n:1614 | C:94.1%[S:33.3%,D:60.7%],<br>F:0.3%,M:5.6%,n:10071 |
| *Pyrus communis* | 1.0 | Hap1 | RAGNAROK | 47,098 | 5,865 | 0.142 | C:97.5%[S:60.7%,D:36.7%],<br>F:1.3%,M:1.2%,n:1614 | C:95.4%[S:32.4%,D:63.0%],<br>F:0.1%,M:4.4%,n:10071 |
| *Pyrus communis* | 1.0 | Hap2 | RAGNAROK | 44,871 | 6,005 | 0.155 | C:97.6%[S:61.3%,D:36.2%],<br>F:1.1%,M:1.3%,n:1614 | C:95.5%[S:33.1%,D:62.4%],<br>F:0.2%,M:4.3%,n:10071 |

**Figure 2.**
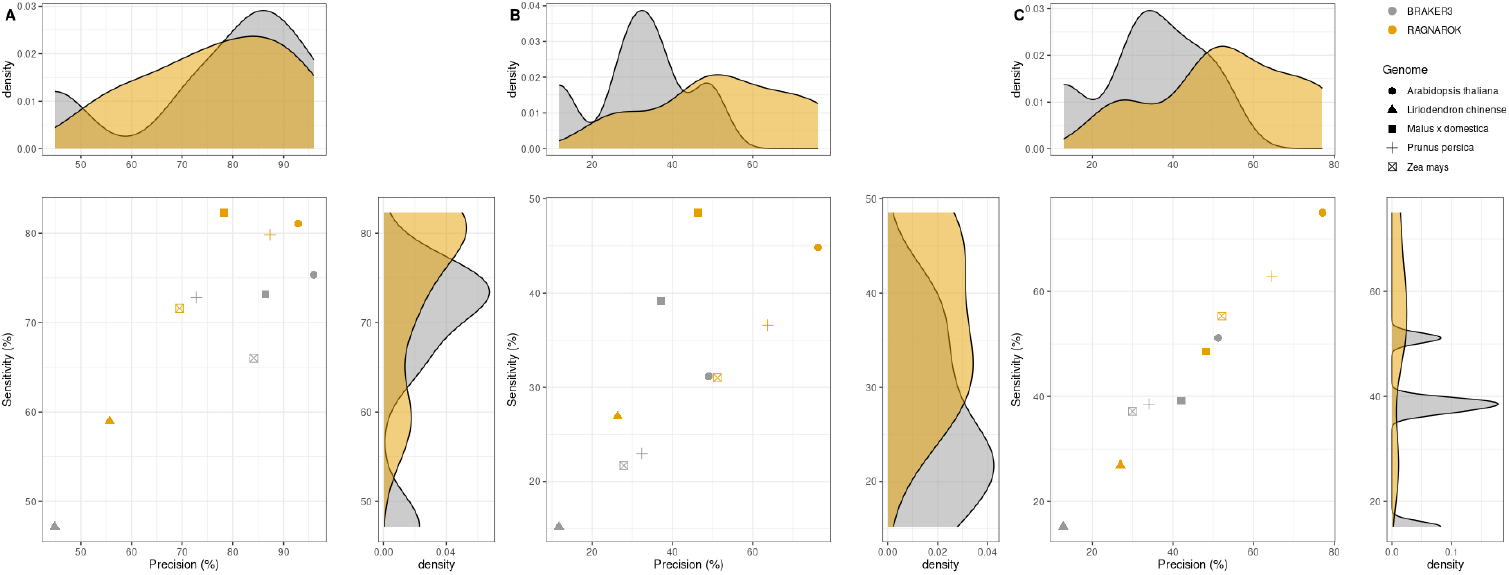
**Precision and sensitivity at different levels of annotation from BRAKER3 and RAGNAROK compared to reference annotation for five genomes. A) Exon level, B) Transcript level, C) Gene level**

We also compared the resulting RAGNAROK, BRAKER3, and reference annotations for each of the five species for the total number of genes annotated, mono-exonic, and mono:multi-exonic rates (Table 1; Fig. 4). Here, neither RAGNAROK nor BRAKER3 produced significantly different annotated total genes compared to the reference. However, total monoexonic genes and the ratio of mono:multi-exonic annotated genes were found to be significant (ANOVA *p-*value = 0.00169 and 0.000817, respectively). In monoexonic gene annotations, RAGNAROK generally exhibited similar numbers to the reference annotation. In contrast, BRAKER3 was significantly higher compared to RAGANROK (*p-*value = 0.00228) and the reference (*p-*value = 0.00472). As a result, the ratio of mono:multi-exonic annotated genes was also significantly different for BRAKER3 compared to RAGNAROK (*p-*value = 0.00108) and the reference annotations (*p-*value = 0.00258). Furthermore, RAGNAROK consistently had a lower number of exons and genes missed compared to BRAKER3. With the exception being BRAKER3 had lower missed genes than RAGNAROK for *Z. mays*. However, the differences were only significant at the exon level (*p-*value = 0.00725). BRAKER3 also exhibited higher numbers of novel genes annotated; however, the difference was not significant (Table 1; Fig. 4).

**Figure 3.**
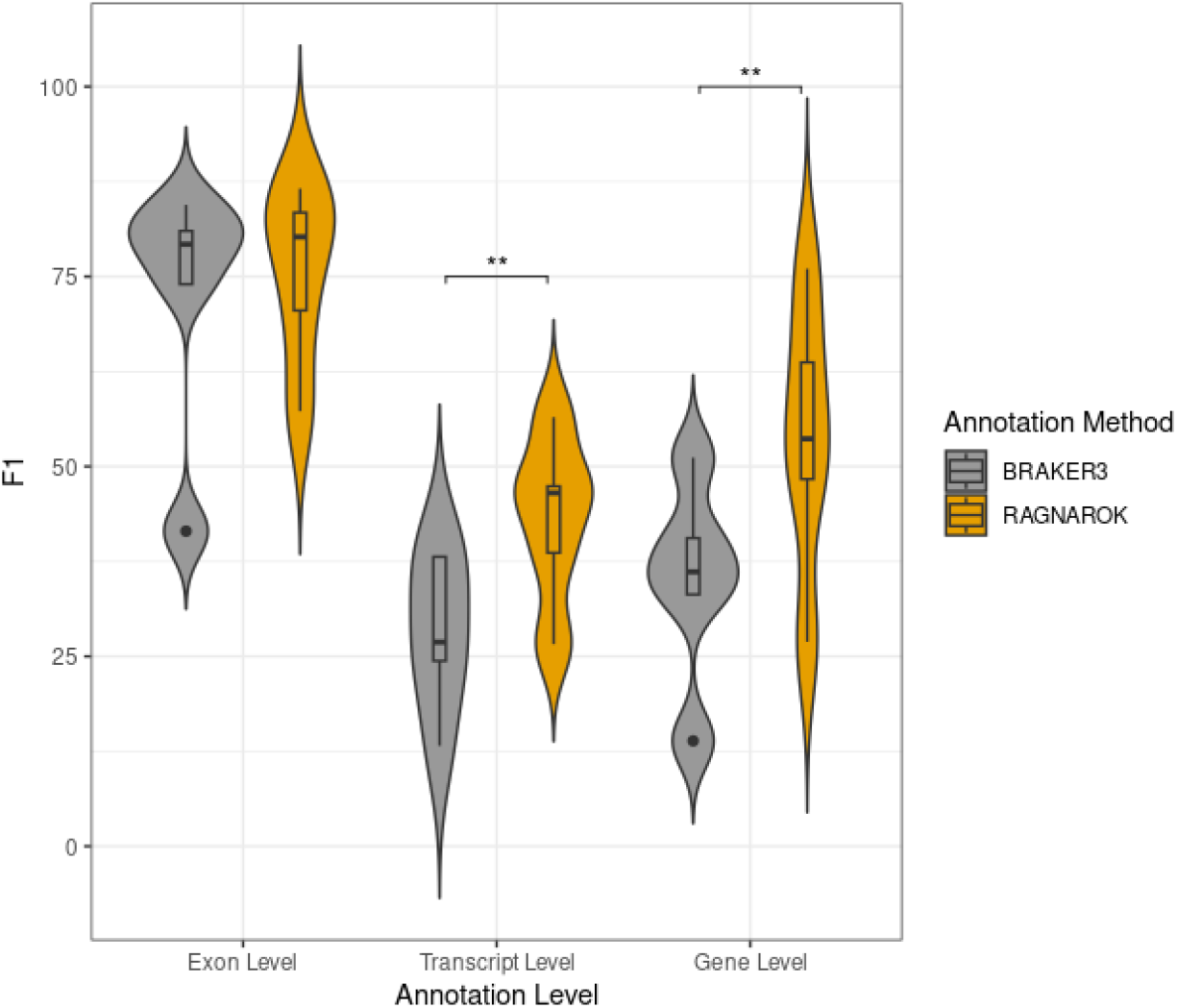
F1 scores between BRAKER3 and RAGNAROK when compared to the reference annotation for five genomes. ^**^ denotes significant differences between the F1 score distributions using a paired t-test and Bonferroni adjustment.

**Figure 4.**
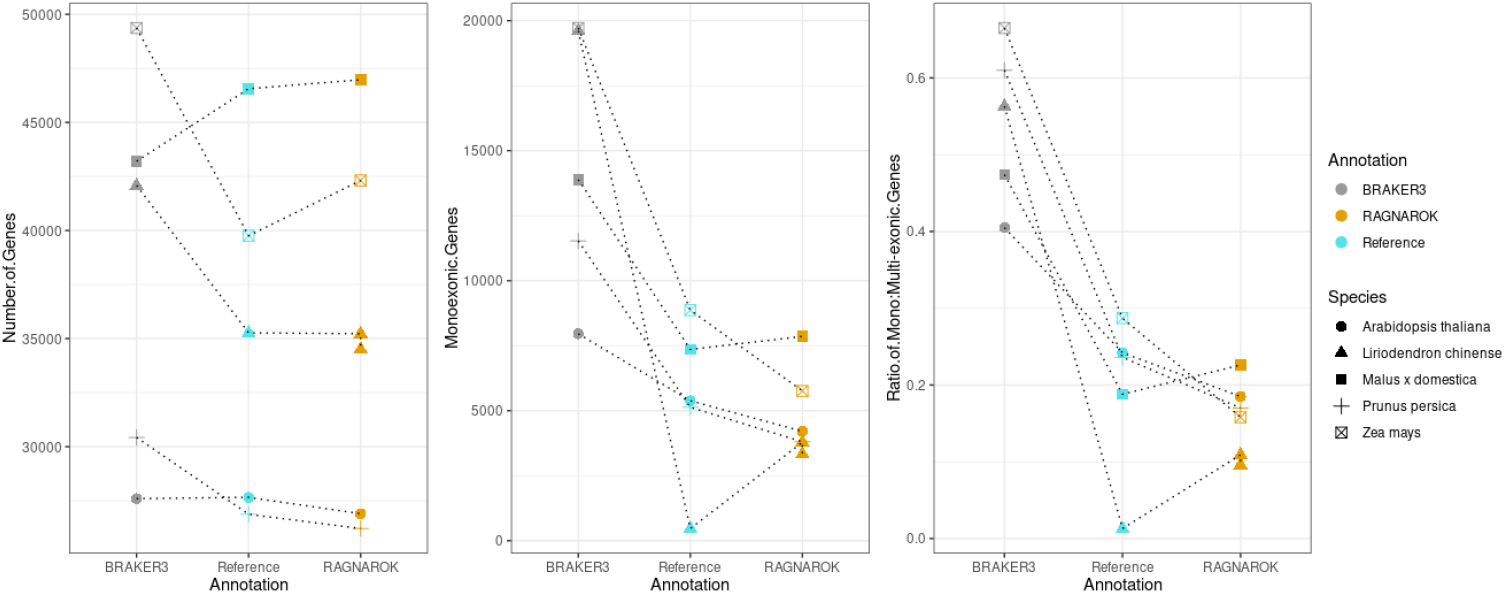
Annotation metrics between BRAKER3 and RAGANAROK compared to the reference annotation for five genomes. Dotted lines connect the paired datasets.

To further evaluate the performance of RAGNAROK compared to BRAKER3 beyond statistical metrics, we compared BUSCO values of the resulting transcriptomes from each annotation method to the reference (Table 1; Fig. 5). BRAKER3 and RAGNAROK exhibited lower BUSCO complete scores compared to the reference annotation for *A. thaliana, P. persica*, and *Z. mays*. In all cases except *A. thaliana* and *Z. mays*, RAGNAROK was higher in the BUSCO complete and fragmented scores over BRAKER3. This trend was inversely consistent with the BUSCO missing category, with RAGNAROK being lower than BRAKER3 except for *Z. mays* (Table 1; Fig. 5).

**Figure 5.**
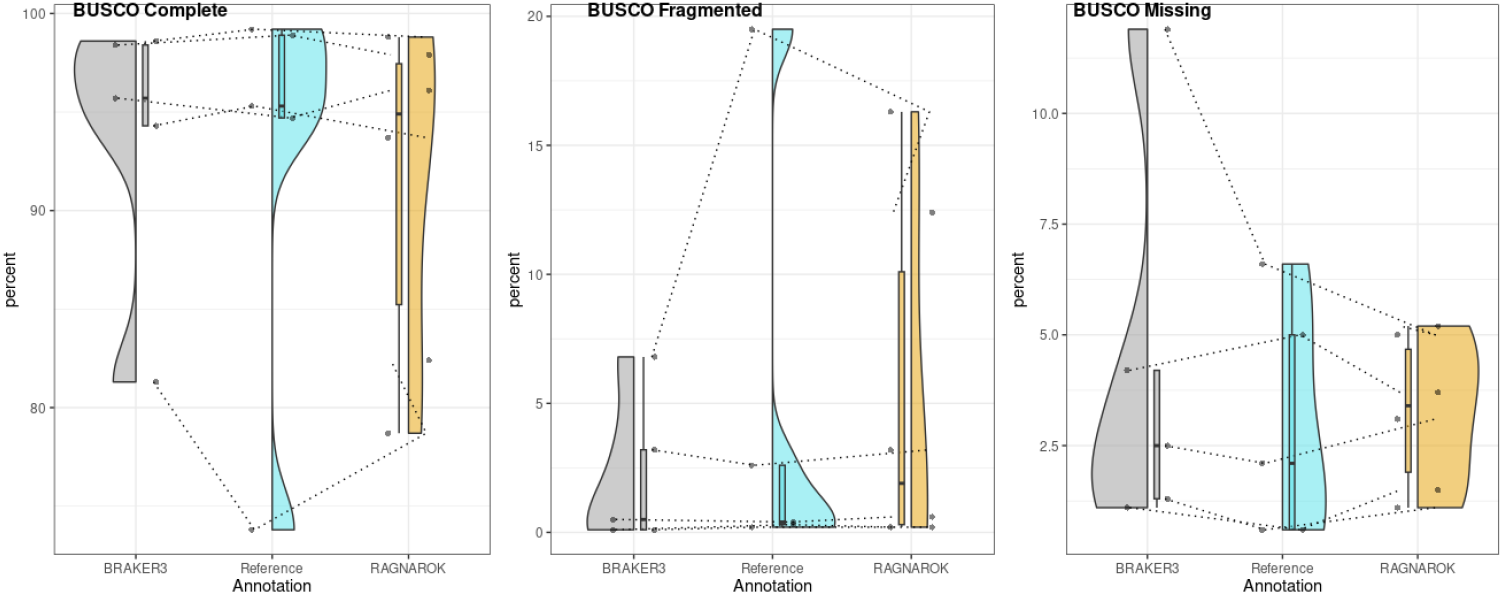
Comparisons of BUSCO scores of BRAKER3, RAGNAROK, and the reference annotations. Dotted lines connect the paired datasets.

### Improvement in reannotation of MAKER-annotated genomes

Our group and others have published and/or released several MAKER-generated annotations for Rosaceae crops (Gottschalk *et al*., 2025; Mansfeld, Ou, *et al*., 2023; Mansfeld, Yocca, *et al*., 2023; Yocca *et al*., 2024). We took this opportunity to re-annotate these five genome assemblies using RAGNAROK to compare against MAKER annotations (Table 2). There was no trend with either annotation method yielding more or fewer genes consistently than the other. However, RAGNAROK annotated significantly higher mono-exonic genes compared to MAKER (*p-*value =0.00781). The resulting ratio of mono:multi-exonic genes was significantly lower in the MAKER annotations compared to RAGNAROK (*p-*value=0.00992) (Table 2). When comparing BUSCO scores using either the general Embryophyta (OBD10) or Rosaceae (OBD12) specific databases, RAGNAROK consistently outperformed MAKER (Fig. 6). RAGNAROK produced significantly higher BUSCO complete, as well as significantly lower fragmented and missing BUSCO genes (Fig. 6).

**Figure 6.**
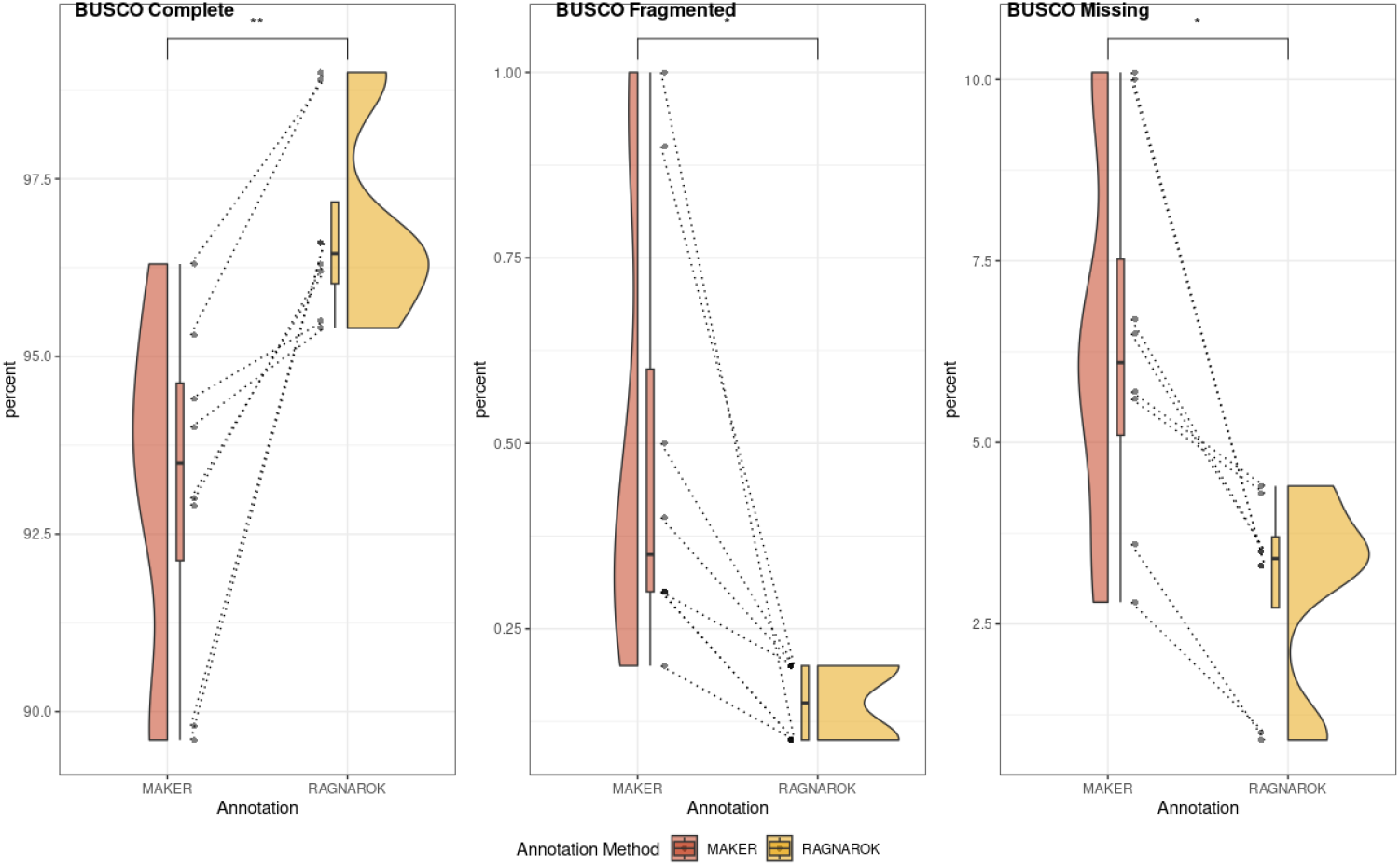
Comparisons of the improvements when reannotating genomes from MAKER to RAGNAROK. Dotted lines connect the paired datasets. ^*^ and ^**^ denote significant differences in means using a paired t-test.

### Run-time comparisons

We recorded the run times for annotating each dataset with BRAKER3 and RAGNAROK. For comparisons, we did not consider the time allocated to TE annotation via EDTA in RAGNAROK to provide a comparable comparison with BRAKER3, which does not include a TE annotation feature (Table 3). Overall, RAGNAROK was faster than BRAKER3, regardless of genome size (Fig. 7). When comparing the regression run time models (Fig. 7), we found the interaction between annotation method and genome size to be significant (ANOVA *p-*value = 0.0268); both BRAKER3 and RAGNAROK run times scaled linearly. The slope of the regression model for RAGNAROK was significantly lower than BRAKER3 (*p-*value *=* 0.0268) (Fig. 7).

**Table 3.** Run-time comparisons between RAGNAROK and BRAKER3.

| Species | Annotation | Repeat Annotation Time |  |  | Gene Annotation Time |  |  |
| --- | --- | --- | --- | --- | --- | --- | --- |
|  |  | Days | Hours | Minutes | Days | Hours | Minutes |
| *Arabidopsis thaliana* | BRAKER3 |  |  |  |  | 4 | 58 |
| *Arabidopsis thaliana* | RAGNAROK |  | 5 | 1 |  | 4 | 22 |
| *Prunus persica* | BRAKER3 |  |  |  |  | 5 | 54 |
| *Prunus persica* | RAGNAROK |  | 9 | 44 |  | 3 | 37 |
| *Malus x domestica* | BRAKER3 |  |  |  |  | 13 | 39 |
| *Malus x domestica* | RAGNAROK | 1 | 5 | 40 |  | 7 | 46 |
| *Liriodendron chinense* | BRAKER3 (short-reads) | EDTA ran outside of RAGNAROK |  |  | 1 | 1 | 30 |
| *Liriodendron chinense* | RAGNAROK (short-reads) |  |  |  |  | 14 | 23 |
| *Liriodendron chinense* | RAGNAROK (long-reads) |  |  |  |  | 15 | 14 |
| *Zea mays* | BRAKER3 |  |  |  | 1 | 2 | 9 |
| *Zea mays* | RAGNAROK | 1 | 17 | 36 |  | 6 | 21 |

| Difference in Gene Annotation Times |  |
| --- | --- |
| *Arabidopsis thaliana* | 36 minutes |
| *Prunus persica* | 2 hours and 17 minutes |
| *Malus x domestica* | 5 hours and 53 minutes |
| *Liriodendron chinense* (short-read) | 11 hours and 7 minutes |
| *Liriodendron chinense* (long-read) | 10 hours and 16 minutes |
| *Zea mays* | 19 hours and 48 minutes |

**Figure 7.**
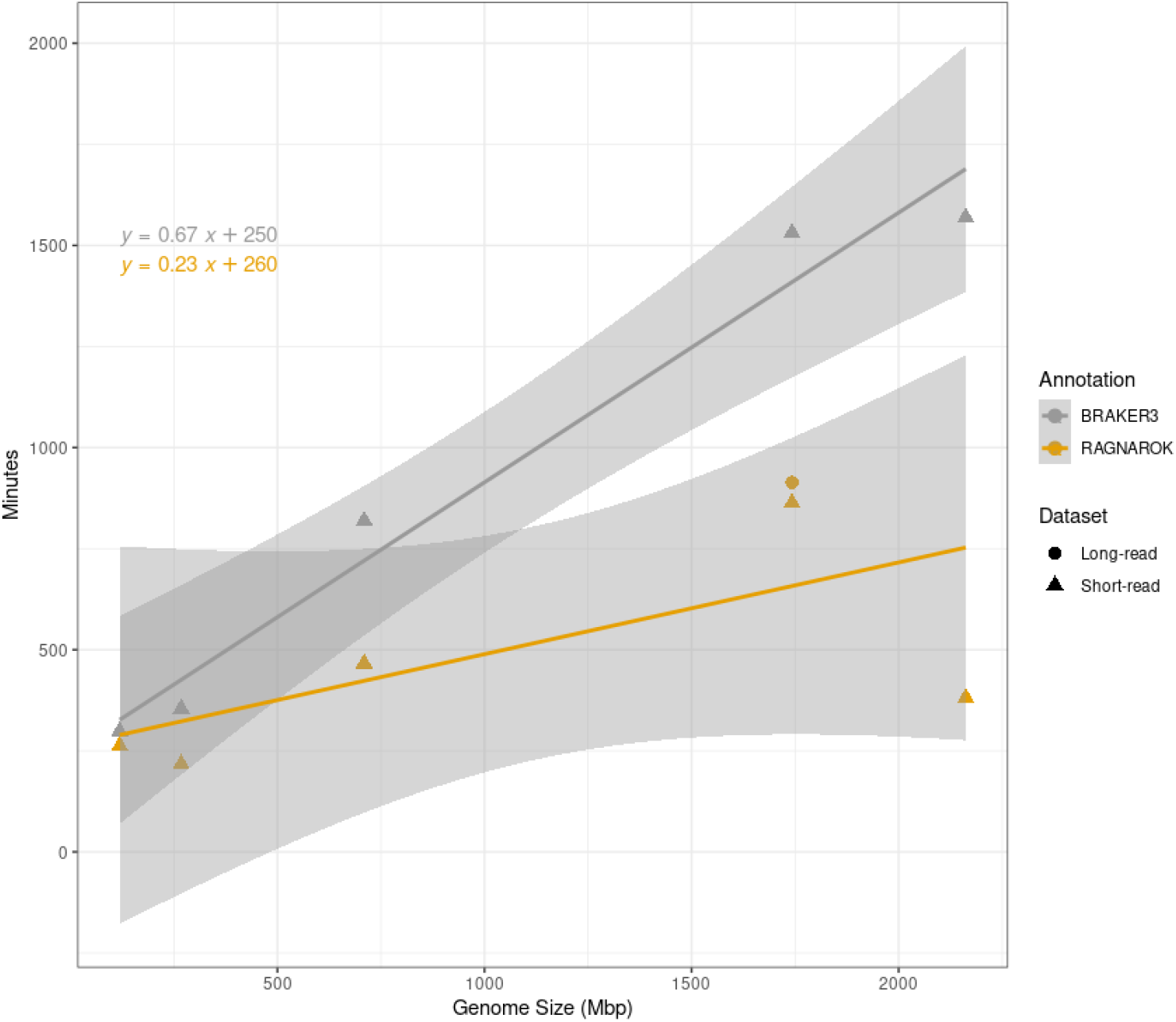
Comparing the run times between BRAKER3 and RAGNAROK annotation pipelines on five datasets. Regression lines were determined using a linear model.

## Discussion

### Installation

We aimed to develop an annotation pipeline that is easy to operate and install. Several other annotation pipelines, such as BRAKER3 and Funannotate, offer a user-friendly installation method using Docker or Apptainer images (Palmer and Stajich, 2020; Gabriel, Brůna, *et al*., 2024; Kurtzer *et al*., 2017; Merkel, 2014). However, both of these pipelines require users to manually download and install the GeneMark predictor, which requires a license agreement and continually updated permission keys to operate (Brůna *et al*., 2024). MAKER requires local installation of the pipeline and all of its dependencies (Campbell *et al*., 2014). To overcome these hurdles and allow for greater accessibility, we elected to rely on Nextflow and Apptainer to manage operations and virtual installation of the dependencies for RAGNAROK (Langer *et al*., 2025; Kurtzer *et al*., 2017). To install and run RAGNAROK, a user will only be required to have Nextflow and Apptainer installed on their system. Furthermore, Nextflow facilitates ease of scalability, which allows RAGNAROK to be run on local machines or on a high-performance computing center (HPCC) (Langer *et al*., 2025).

### Performance against other annotation pipelines

To compare performance against BRAKER3, the current top annotation pipeline, we annotated five reference-quality genomes (Gabriel, Brůna, *et al*., 2024). These genomes include the small genome model plant *Arabidopsis*, as well as larger fruit crops of peach and apple, and lastly, Gb-scale genomes of maize and *L. chinense* (Table 1) (Chen *et al*., 2019; Lamesch *et al*., 2012; Verde *et al*., 2013, 2017; Daccord *et al*., 2017; Hufford *et al*., 2021). We undertook a similar approach as the BRAKER3 authors by limiting the amount of input data to reflect real-world conditions (Gabriel, Brůna, *et al*., 2024). We found that RAGNAROK produced a higher level of exon, transcript, and gene-level annotation accuracy compared to BRAKER3 (Fig. 2). In most annotation levels (exon, transcript, gene), RAGNAROK was significantly higher (STable 2). As a result, RAGNAROK produced significantly higher transcript and gene F1 scores (STable 2). Furthermore, RAGNAROK demonstrated lower monoexonic gene annotations than BRAKER3 (Table 2; Fig. 4). Other authors have found similar high rates of monoexonic gene annotations when using BRAKER (Venturini *et al*., 2018). We observed mono:multi-exonic ratios of ∼0.17 with RAGNAROK, which is close to the suggested ideal ratio of 0.2 (Venturini *et al*., 2018; Jain *et al*., 2008). Several of the reference annotations used for benchmarking were previously generated using BRAKER3, which may have contributed to the observed concordance in gene model predictions in maize and the general trend of mono-exonic:multi-exonic ratios greater than 0.2 in the reference (Table 1). In all cases except *A. thaliana* and *Z. mays*, RAGNAROK was higher in the BUSCO complete and fragmented scores over BRAKER3 (Table 1; Fig. 5). This result could be attributed to methodological consistency of these annotations with those found in the BUSCO lineage databases, including monoexonic genes. This will require further analysis to conclude. BRAKER3 and RAGNAROK exhibited lower BUSCO complete scores compared to the reference annotation for *A. thaliana, P. persica, and Z. mays*. However, the differences were minimal, with the maximum difference in Complete Gene percentage being <2% (Table 1). This is not a surprising result, as all three of these species have undergone manual curation to their annotation and utilized more evidential datasets during their annotation (Lamesch *et al*., 2012; Verde *et al*., 2013, 2017; Hufford *et al*., 2021).

We also compared RAGANROK to MAKER2 annotations for a set of Rosaceae genomes (Gottschalk *et al*., 2025; Mansfeld, Yocca, *et al*., 2023; Mansfeld, Ou, *et al*., 2023; Yocca *et al*., 2024). Since these were not reference genome annotations, sensitivity, precision, and F1 scores could not be assessed. However, we did carry out comparisons on annotated gene number, monoexonic annotations, ratios, and BUSCO. In contrast to the comparisons with BRAKER, MAKER2 produced significantly fewer monoexonic gene annotations than RAGANROK. However, the average ratio for MAKER2 (∼0.11) was below the suggested ratio of 0.2 (Venturini *et al*., 2018; Jain *et al*., 2008). Whereas, RAGNAROK maintained a ratio of ∼0.19. When comparing BUSCO scores between MAKER2 and RAGNAROK, RAGNAROK consistently outperformed MAKER2 (Fig. 6; Table 2). This result was demonstrated using the >10k Rosaceae-specific ODB12 BUSCO dataset (Tegenfeldt *et al*., 2025). Here, RAGNAROK achieved a complete score averaging ∼96.8% compared to 93.2% for MAKER2, a significant increase (Fig. 6).

### Run-time

RAGNAROK demonstrated consistent and significantly faster run times across five plant genomes of various lengths compared to BRAKER3 (Fig. 7). We observed slightly faster BRAKER3 runtimes than what was demonstrated in the BRAKER3 publication (Gabriel, Brůna, *et al*., 2024). For example, that publication reported a BRAKER3 runtime of 5h 37m for Arabidopsis compared to the 4h 58m report in this study (Gabriel, Brůna, *et al*., 2024). This improvement could be associated with revisions to the BRAKER3 pipeline since its release. However, the resource allocation between BRAKER3 and RAGNAROK can conflate the performance in favor of RAGNAROK. For example, BRAKER3 is thread-limited to 48 threads due to a max level in the optimize_augustus.pl script. In contrast, RAGNAROK can be scaled to whatever is available on the local system or the HPCC, and some of the subprocesses (*i*.*e*., Helixer) are GPU-accelerated.

### Future Directions

We aim to introduce further improvements and broaden the usage of RAGNAROK in future versions. Currently, we have limited RAGNAROK to plants, but anticipate adding flexibility for other Eukaryotes such as animals and fungi. This will be achieved through the implementation of user-selected specific Helixer models released by the software’s developers (Stiehler *et al*., 2021; Holst *et al*., 2023). There are further opportunities to allow user-generated Helixer models trained on RNA-seq data. Similarly, as other novel tools and models become available, we envision adding support for additional annotation software (*e*.*g*., Tiberius) (Gabriel, Becker, *et al*., 2024) to continue to leverage the advantages of both traditional and deep-learning-based approaches. The modular implementation of RAGNAROK and the parallel processing efficiency of Nextflow will continue to support the ongoing development and expansion of pipeline features. Nevertheless, the current implementation of RAGNAROK is already capable of incorporating gene models from such emerging tools with the addition of manual weighting from the end-user.

We also plan to incorporate additional subparameters to enhance user-specified intron length during the RNA-seq alignment step performed by STAR (Dobin *et al*., 2013). Currently, it is encoded at a 10Kb max intron size to support what is commonly observed in plants. However, allowing flexibility will be required for use with other Eukaryotes. For example, when annotating a fungal genome, the max intron length should be reduced from the current 10kb to ∼2kb (Kupfer *et al*., 2004). We also plan to integrate the FindPlantNLR pipeline as an additional annotation process (Chen *et al*., 2023b). Our team has found FindPlantNLR offers improvements in annotating complex nucleotide-binding domain (NBD) leucine-rich repeat (LRR) proteins (NLRs), which are commonly found in complex tandem duplications (Larson *et al*., 2025; Liu *et al*., 2025). These annotation tools will further serve in preserving annotations when conducting reannotation of genomes. We also envision some improvements to decrease run time. For example, there are other emerging GPU-accelerated implementations of standard bioinformatic pipelines through programs such as Parabricks from NVIDIA (Santa Clara, CA)(Franke and Crowgey, 2020). Tasks such as RNAseq mapping using STAR and minimap have already been made available through Parabricks and would be complementary to the GPU-accelerated Helixer gene prediction. Lastly, we expect to integrate hybrid transcript assembly from mixed short-read and long-read data as implemented in StringTie2 (Shumate *et al*., 2022).

## Availability

To download and utilize the RAGNAROK pipeline, please refer to its repository on GitHub at the following web address: https://github.com/ryandkuster/ragnarok.

## Contributions

CG conceptualized the study and developed the initial methodology. RK advanced the methodology, implementation using NextFlow, and maintains the pipeline. ZS, LW, MS, and BNM contributed methodology improvements, testing, and validation of the pipeline. All authors contributed to the writing, editing, and reviewing of the manuscript.

## Acknowledgments

We appreciate input and feedback from Dr. Chris Dardick and Shade Niece during the development of the pipeline and evaluation of its performance.

## Supplemental

Supplemental Table 1. Genomes, reference annotations, evidence datasets, and versions for different species used in the comparisons between annotation pipelines.

Supplemental Table 2. Precision and sensitivity for gene predictions by RAGNAROK and BRAKER3 compared to the reference annotation for five reference quality genomes.

Supplemental File 1. Commands executed in the comparisons between RAGNAROK, BRAKER3, and MAKER.

