## Supplemental File 1 for "Ragnarok: a flexible and RApid GeNe Annotation (ROcKs) pipeline deployed through Nextflow"

**Supplemental File 1. Script log for benchmarking of RAGNAROK vs BRAKER3.**

A) *Arabidopsis thaliana*

```
#!/bin/bash

mkdir rna
cd rna

#download the SRA for RNAseq evidence
prefetch SRR8714016
prefetch SRR8759751
prefetch SRR4010853
prefetch SRR7289569
prefetch SRR12547664
prefetch SRR12076896

fasterq-dump -S SRR8714016
fasterq-dump -S SRR8759751
fasterq-dump -S SRR4010853
fasterq-dump -S SRR7289569
fasterq-dump -S SRR12547664
fasterq-dump -S SRR12076896

cd ..

############################################################
#                        Ragnarok!                         #
############################################################

nextflow  ~/Programs/ragnarok-main/main.nf \
   --publish_dir    ${PWD}
   --genome        ${PWD}/Athaliana_447_TAIR10.fa \
   --cds          ${PWD}/Athaliana_447_Araport11.cds_primaryTranscriptOnly.fa \
   --protein       ${PWD}/Athaliana_447_Araport11.protein_primaryTranscriptOnly.fa
\
   --ill          ${PWD}/rna \
   --perform_masking true \
   --skip_qc       false \
   --skip_trim      false \
   --nlrs          false \
   --design       ~/Programs/ragnarok-main/assets/mikado_conf.tsv \
   --scoring      ~/Programs/ragnarok-main/plant.yaml \
   --homology     ${PWD}/uniprot_sprot.fasta \
   --contam        "insecta,fungi,bacteria" \
   -profile local,eight \
```

#copy the Athaliana_447_TAIR10.fa.mod.MAKER.masked.fa from the EDTA output within the /work/ directory generated by RAGNAROK. It will be used as the masked input into BRAKER3

```
cp ./work/path/to/job/id/Athaliana_447_TAIR10.fa.mod.MAKER.masked.fa ./

###########################################################
#                          BRAKER3                        #
###########################################################

singularity exec -B ${PWD}:${PWD} ${BRAKER_SIF} braker.pl --threads 48 --
workingdir=${PWD} \
--genome=${PWD}/Athaliana_447_TAIR10.fa.mod.MAKER.masked --
species=arabidopsis_new \
--prot_seq=${PWD}/Athaliana_447_Araport11.protein_primaryTranscriptOnly.fa \
  --
rnaseq_sets_ids=SRR8714016,SRR8759751,SRR4010853,SRR7289569,SRR12547664,SRR
12076896 \
--rnaseq_sets_dirs=${PWD}/RNA/

###########################################################
#                        comparisons                      #
###########################################################

mkdir metrics
cd metrics

mikado util stats ${PWD}/Athaliana_447_Araport11.gene.gff3
Athaliana_447_Araport11.gene.stats

~/Augustus/scripts/gtf2gff.pl <${PWD}/braker3_anno/braker.gtf --gff3 --
out=/${PWD}/braker.gff

mikado util stats ${PWD}/braker.gff braker.stats

mikado util stats ${PWD}/publish/entap_final/mikado.loci_out.entap_filtered.gff3
ragnarok.stats

mikado compare -r Athaliana_447_Araport11.gene.gff3 -p braker.gff -o
braker.metrics -x 30

mikado compare -r Athaliana_447_Araport11.gene.gff3 -p
${PWD}/publish/entap_final/mikado.loci_out.entap_filtered.gff3 -o
ragnarok.metrics -x 30
```

B) *Prunus persica*
*#downloaded Prunus persica v2.0.a1 annotation and assembly*

```
wget
https://www.rosaceae.org/rosaceae_downloads/Prunus_persica/Prunus_persica-genome.v2.0.a1/assembly/Prunus_persica_v2.0.a1_scaffolds.fasta.gz
wget
https://www.rosaceae.org/rosaceae_downloads/Prunus_persica/Prunus_persica-genome.v2.0.a1/genes/Prunus_persica_v2.0.a1.gene.gff3.gz
wget
https://www.rosaceae.org/rosaceae_downloads/Prunus_persica/Prunus_persica-genome.v2.0.a1/genes/Prunus_persica_v2.0.a1.primaryTrs.pep.fa.gz
wget
https://www.rosaceae.org/rosaceae_downloads/Prunus_persica/Prunus_persica-genome.v2.0.a1/genes/Prunus_persica_v2.0.a1.primaryTrs.cds.fa.gz

#download the SRAs used in the v1 genome paper
mkdir rna
cd rna

prefetch SRR531865
prefetch SRR531863
prefetch SRR531864
prefetch SRR531862

fasterq-dump -m 25GB -e 90 -p -S SRR531865
fasterq-dump -m 25GB -e 90 -p -S SRR531863
fasterq-dump -m 25GB -e 90 -p -S SRR531864
fasterq-dump -m 25GB -e 90 -p -S SRR531862

cd ..

###########################################################
#                        Ragnarok!                        #
###########################################################
nextflow  ~/Programs/ragnarok-main/main.nf \
   --publish_dir     ${PWD}
   --genome        ${PWD}/Prunus_persica_v2.0.a1.scaffolds.fasta \
   --cds           ${PWD}/Prunus_persica_v2.0.a1.primaryTrs.cds.fa \
   --protein       ${PWD}/Prunus_persica_v2.0.a1.primaryTrs.pep.fa \
   --ill           ${PWD}/rna \
   --perform_masking true \
   --skip_qc       false \
   --skip_trim     false \
   --nlrs          false \
   --design        ~/Programs/ragnarok-main/assets/mikado_conf.tsv \
   --scoring       ~/Programs/ragnarok-main/plant.yaml \
   --homology      ${PWD}/uniprot_sprot.fasta \
```

```
  --contam        "insecta,fungi,bacteria" \
  -profile local,eight \

############################################################
#                          BRAKER3                         #
############################################################

singularity exec -B ${PWD}:${PWD} ${BRAKER_SIF} braker.pl --threads 48 --
workingdir=${PWD} \
--genome=${PWD}/Prunus_persica_v2.0.a1.scaffolds.fasta.mod.MAKER.masked --
species=peach \
--prot_seq=${PWD}/Prunus_persica_v2.0.a1.primaryTrs.pep.fa \
--rnaseq_sets_ids=SRR531865,SRR531863,SRR531864,SRR531862 \
--rnaseq_sets_dirs=${PWD}/rna/

############################################################
#                        comparisons                       #
############################################################

mkdir metrics
cd metrics

mikado util stats ${PWD}/Prunus_persica_v2.0.a1.gene.gff3
Prunus_persica_v2.0.a1.gene.stats

~/Augustus/scripts/gtf2gff.pl <${PWD}/braker3_anno/braker.gtf --gff3  --
out=/${PWD}/braker.gff

mikado util stats ${PWD}/braker.gff braker.stats

mikado util stats ${PWD}/publish/entap_final/mikado.loci_out.entap_filtered.gff3
ragnarok.stats

mikado compare -r Prunus_persica_v2.0.a1.gene.gff3 -p braker.gff -o braker.metrics
-x 30

mikado compare -r Prunus_persica_v2.0.a1.gene.gff3 -p
${PWD}/publish/entap_final/mikado.loci_out.entap_filtered.gff3 -o
ragnarok.metrics -x 30
```

C) *Malus domestica*
*#downloaded GDDH13 v1.1 annotation and assembly*

```
wget
https://www.rosaceae.org/rosaceae_downloads/Malus_x_domestica/Malus_x_domestica-genome_GDDH13_v1.1/assembly/GDDH13_1-1_formatted.fasta.gz
wget
https://www.rosaceae.org/rosaceae_downloads/Malus_x_domestica/Malus_x_domestica-genome_GDDH13_v1.1/genes/gene_models_20170612.gff3.gz
wget
https://www.rosaceae.org/rosaceae_downloads/Malus_x_domestica/Malus_x_domestica-genome_GDDH13_v1.1/genes/GDDH13_1-1_prot.fasta.gz
wget
https://www.rosaceae.org/rosaceae_downloads/Malus_x_domestica/Malus_x_domestica-genome_GDDH13_v1.1/genes/GDDH13_1-1_mrna.fasta.gz


#download the SRAs used in the v1 genome paper
mkdir rna
cd rna

prefetch SRR767660
prefetch SRR767668
prefetch SRR767669
prefetch SRR767670
prefetch SRR767671
prefetch SRR767672
prefetch SRR767673
prefetch SRR767674
prefetch SRR768127
prefetch SRR768128
prefetch SRR768129
prefetch SRR768130
prefetch SRR768131
prefetch SRR768132
prefetch SRR768133
prefetch SRR768134
prefetch SRR768135
prefetch SRR768136
prefetch SRR768137


fasterq-dump -m 25GB -e 90 -p -S SRR767660
fasterq-dump -m 25GB -e 90 -p -S SRR767668
fasterq-dump -m 25GB -e 90 -p -S SRR767669
fasterq-dump -m 25GB -e 90 -p -S SRR767670
fasterq-dump -m 25GB -e 90 -p -S SRR767671
fasterq-dump -m 25GB -e 90 -p -S SRR767672
```

```
fasterq-dump -m 25GB -e 90 -p -S SRR767673
fasterq-dump -m 25GB -e 90 -p -S SRR767674
fasterq-dump -m 25GB -e 90 -p -S SRR768127
fasterq-dump -m 25GB -e 90 -p -S SRR768128
fasterq-dump -m 25GB -e 90 -p -S SRR768129
fasterq-dump -m 25GB -e 90 -p -S SRR768130
fasterq-dump -m 25GB -e 90 -p -S SRR768131
fasterq-dump -m 25GB -e 90 -p -S SRR768132
fasterq-dump -m 25GB -e 90 -p -S SRR768133
fasterq-dump -m 25GB -e 90 -p -S SRR768134
fasterq-dump -m 25GB -e 90 -p -S SRR768135
fasterq-dump -m 25GB -e 90 -p -S SRR768136
fasterq-dump -m 25GB -e 90 -p -S SRR768137


cd ..

############################################################
#                          Ragnarok!                       #
############################################################
nextflow  ~/Programs/ragnarok-main/main.nf \
   --publish_dir     ${PWD}
   --genome         ${PWD}/GDDH13_1-1_formatted.fasta \
   --cds            ${PWD}/GDDH13_1-1_mrna.fasta \
   --protein        ${PWD}/GDDH13_1-1_prot.fasta \
   --ill            ${PWD}/rna \
   --perform_masking true \
   --skip_qc        false \
   --skip_trim      false \
   --nlrs           false \
   --design         ~/Programs/ragnarok-main/assets/mikado_conf.tsv \
   --scoring        ~/Programs/ragnarok-main/plant.yaml \
   --homology       ${PWD}/uniprot_sprot.fasta \
   --contam         "insecta,fungi,bacteria" \
   -profile local,eight \

############################################################
#                          BRAKER3                         #
############################################################

singularity exec -B ${PWD}:${PWD} ${BRAKER_SIF} braker.pl --threads 48 --
workingdir=${PWD} \
--genome=${PWD}/GDDH13_1-1_formatted.fasta.mod.MAKER.masked --
species=GDDH13 \
--prot_seq=${PWD}/GDDH13_1-1_prot.fasta \
--
rnaseq_sets_ids=SRR767660,SRR767668,SRR767669,SRR767670,SRR767671,SRR767672
```

```
,SRR767673,SRR767674,SRR768127,SRR768132,SRR768133,SRR768134,SRR768135,SRR768136,SRR768137 \
--rnaseq_sets_dirs=${PWD}/rna/

############################################################
#                        comparisons                       #
############################################################

mkdir metrics
cd metrics

mikado util stats ${PWD}/gene_models_20170612.gff3 GDDH13.gene.stats

~/Augustus/scripts/gtf2gff.pl <${PWD}/braker3_anno/braker.gtf --gff3 --out=/${PWD}/braker.gff

mikado util stats ${PWD}/braker.gff braker.stats

mikado util stats ${PWD}/publish/entap_final/mikado.loci_out.entap_filtered.gff3 ragnarok.stats

mikado compare -r gene_models_20170612.gff3 -p braker.gff -o braker.metrics -x 30

mikado compare -r gene_models_20170612.gff3 -p ${PWD}/publish/entap_final/mikado.loci_out.entap_filtered.gff3 -o ragnarok.metrics -x 30
```

D) *Liriodendron chinense*

```
#downloaded Lich 1.0 annotation and assembly

wget https://treegenesdb.org/FTP/Genomes/Lich/v1.0/genome/Lich.1_0.fa
wget https://treegenesdb.org/FTP/Genomes/Lich/v1.0/annotation/Lich.1_0.gff
wget https://treegenesdb.org/FTP/Genomes/Lich/v1.0/annotation/Lich.1_0.pep.fa.gz
wget https://treegenesdb.org/FTP/Genomes/Lich/v1.0/annotation/Lich.1_0.cds.fa.gz

#had to run EDTA separately due to it crashing the system (needed more threads and memory)

edta.pl --genome Lich.1_0.fa --cds Lich.1_0.cds.fa --overwrite 1 --sensitive 1 --anno 1 --force 1 --threads 96

#download the SRAs used in the v1 genome paper
mkdir rna/ill
cd rna/ill

#illumina
```

```
prefetch SRR9945429
prefetch SRR9945430
prefetch SRR9945431
prefetch SRR9945432
prefetch SRR9945433
prefetch SRR9945434
prefetch SRR9945435
prefetch SRR9948913
prefetch SRR9948914
prefetch SRR9948915
prefetch SRR9948916
prefetch SRR9948917
prefetch SRR9948918
prefetch SRR9948919
prefetch SRR9949005
prefetch SRR9949006
prefetch SRR9949007
prefetch SRR9949008
prefetch SRR9949009
prefetch SRR9949010
prefetch SRR9949011

fasterq-dump -m 25GB -e 90 -p -S SRR9945429
fasterq-dump -m 25GB -e 90 -p -S SRR9945430
fasterq-dump -m 25GB -e 90 -p -S SRR9945431
fasterq-dump -m 25GB -e 90 -p -S SRR9945432
fasterq-dump -m 25GB -e 90 -p -S SRR9945433
fasterq-dump -m 25GB -e 90 -p -S SRR9945434
fasterq-dump -m 25GB -e 90 -p -S SRR9945435
fasterq-dump -m 25GB -e 90 -p -S SRR9948913
fasterq-dump -m 25GB -e 90 -p -S SRR9948914
fasterq-dump -m 25GB -e 90 -p -S SRR9948915
fasterq-dump -m 25GB -e 90 -p -S SRR9948916
fasterq-dump -m 25GB -e 90 -p -S SRR9948917
fasterq-dump -m 25GB -e 90 -p -S SRR9948918
fasterq-dump -m 25GB -e 90 -p -S SRR9948919
fasterq-dump -m 25GB -e 90 -p -S SRR9949005
fasterq-dump -m 25GB -e 90 -p -S SRR9949006
fasterq-dump -m 25GB -e 90 -p -S SRR9949007
fasterq-dump -m 25GB -e 90 -p -S SRR9949008
fasterq-dump -m 25GB -e 90 -p -S SRR9949009
fasterq-dump -m 25GB -e 90 -p -S SRR9949010
fasterq-dump -m 25GB -e 90 -p -S SRR9949011

#iso-seq
mkdir iso
cd iso
```

```
prefetch SRR12695187
fasterq-dump -m 25GB -e 90 -p -SSRR12695187

cd .../..

##########################################################
#                         Ragnarok!                      #
##########################################################
#illumina only annotation
nextflow  ~/Programs/ragnarok-main/main.nf \
   --publish_dir    ${PWD}/ill
   --genome        ${PWD}/Lich.1_0.fa.mod.MAKER.masked \
   --cds           ${PWD}/Lich.1_0.cds.fa \
   --protein       ${PWD}/Lich.1_0.pep.fa \
   --ill           ${PWD}/rna/ill \
   --perform_masking true \
   --skip_qc       false \
   --skip_trim      false \
   --nlrs          false \
   --design        ~/Programs/ragnarok-main/assets/mikado_conf.tsv \
   --scoring      ~/Programs/ragnarok-main/plant.yaml \
   --homology       ${PWD}/uniprot_sprot.fasta \
   --contam        "insecta,fungi,bacteria" \
   -profile local,eight \

#iso-seq only annotation
nextflow  ~/Programs/ragnarok-main/main.nf \
   --publish_dir    ${PWD}/iso
   --genome        ${PWD}/Lich.1_0.fa.mod.MAKER.masked \
   --cds           ${PWD}/Lich.1_0.cds.fa \
   --protein       ${PWD}/Lich.1_0.pep.fa \
   --iso           ${PWD}/rna/iso \
   --perform_masking true \
   --skip_qc       false \
   --skip_trim      false \
   --nlrs          false \
   --design        ~/Programs/ragnarok-main/assets/mikado_conf.tsv \
   --scoring      ~/Programs/ragnarok-main/plant.yaml \
   --homology       ${PWD}/uniprot_sprot.fasta \
   --contam        "insecta,fungi,bacteria" \
   -profile local,eight \

##########################################################
#                         BRAKER3                        #
##########################################################

singularity exec -B ${PWD}:${PWD} ${BRAKER_SIF} braker.pl --threads 48 --
workingdir=${PWD} \
```

```
--genome=${PWD}/Lich.1_0.fa.mod.MAKER.masked --species=Lich \
--prot_seq=${PWD}/Lich.1_0.pep.fa \
--
rnaseq_sets_ids=SRR9945429,SRR9945430,SRR9945431,SRR9945432,SRR9945433,SRR9945434,SRR9945435,SRR9948913,SRR9948914,SRR9948915,SRR9948916,SRR9948917,SRR9948918,SRR9948919,SRR9949005,SRR9949006,SRR9949007,SRR9949008,SRR9949009,SRR9949010,SRR9949011 \
--rnaseq_sets_dirs=${PWD}/rna/ill

###########################################################
#                      comparisons                        #
###########################################################

mkdir metrics
cd metrics

mikado util stats ${PWD}/Lich.1_0.gff3 Lich.1_0.gene.stats

~/Augustus/scripts/gtf2gff.pl <${PWD}/braker3_anno/braker.gtf --gff3 --out=/${PWD}/braker.gff

mikado util stats ${PWD}/braker.gff braker.stats

mikado util stats ${PWD}/publish/entap_final/mikado.loci_out.entap_filtered.gff3 ragnarok.stats

mikado compare -r Lich.1_0.gff3 -p braker.gff -o braker.metrics -x 30

mikado compare -r Lich.1_0.gff3 -p ${PWD}/publish/entap_final/mikado.loci_out.entap_filtered.gff3 -o ragnarok.metrics -x 30
```

E) *Zea mays*

```
#!/usr/bin/env bash
wget https://download.maizegdb.org/Zm-B73-REFERENCE-NAM-5.0/Zm-B73-REFERENCE-NAM-5.0_Zm00001eb.1.canonical.cds.fa.gz
wget https://download.maizegdb.org/Zm-B73-REFERENCE-NAM-5.0/Zm-B73-REFERENCE-NAM-5.0_Zm00001eb.1.protein.fa.gz
wget https://download.maizegdb.org/Zm-B73-REFERENCE-NAM-5.0/Zm-B73-REFERENCE-NAM-5.0.fa.gz
```

```
#!/usr/bin/env bash

wget -nc ftp://ftp.sra.ebi.ac.uk/vol1/fastq/ERR377/003/ERR3773813/ERR3773813_2.fastq.gz
wget -nc ftp://ftp.sra.ebi.ac.uk/vol1/fastq/ERR377/007/ERR3773807/ERR3773807_1.fastq.gz
wget -nc ftp://ftp.sra.ebi.ac.uk/vol1/fastq/ERR377/005/ERR3773825/ERR3773825_1.fastq.gz
```

```
wget -nc ftp://ftp.sra.ebi.ac.uk/vol1/fastq/ERR377/004/ERR3773814/ERR3773814_2.fastq.gz
wget -nc ftp://ftp.sra.ebi.ac.uk/vol1/fastq/ERR377/000/ERR3773810/ERR3773810_1.fastq.gz
wget -nc ftp://ftp.sra.ebi.ac.uk/vol1/fastq/ERR377/007/ERR3773827/ERR3773827_2.fastq.gz
wget -nc ftp://ftp.sra.ebi.ac.uk/vol1/fastq/ERR377/007/ERR3773817/ERR3773817_2.fastq.gz
wget -nc ftp://ftp.sra.ebi.ac.uk/vol1/fastq/ERR377/001/ERR3773821/ERR3773821_1.fastq.gz
wget -nc ftp://ftp.sra.ebi.ac.uk/vol1/fastq/ERR377/006/ERR3773816/ERR3773816_2.fastq.gz
wget -nc ftp://ftp.sra.ebi.ac.uk/vol1/fastq/ERR377/000/ERR3773820/ERR3773820_1.fastq.gz
wget -nc ftp://ftp.sra.ebi.ac.uk/vol1/fastq/ERR377/007/ERR3773827/ERR3773827_1.fastq.gz
wget -nc ftp://ftp.sra.ebi.ac.uk/vol1/fastq/ERR377/001/ERR3773811/ERR3773811_2.fastq.gz
wget -nc ftp://ftp.sra.ebi.ac.uk/vol1/fastq/ERR377/005/ERR3773815/ERR3773815_2.fastq.gz
wget -nc ftp://ftp.sra.ebi.ac.uk/vol1/fastq/ERR377/009/ERR3773809/ERR3773809_1.fastq.gz
wget -nc ftp://ftp.sra.ebi.ac.uk/vol1/fastq/ERR377/002/ERR3773822/ERR3773822_1.fastq.gz
wget -nc ftp://ftp.sra.ebi.ac.uk/vol1/fastq/ERR377/006/ERR3773826/ERR3773826_1.fastq.gz
wget -nc ftp://ftp.sra.ebi.ac.uk/vol1/fastq/ERR377/000/ERR3773810/ERR3773810_2.fastq.gz
wget -nc ftp://ftp.sra.ebi.ac.uk/vol1/fastq/ERR377/002/ERR3773812/ERR3773812_2.fastq.gz
wget -nc ftp://ftp.sra.ebi.ac.uk/vol1/fastq/ERR377/004/ERR3773824/ERR3773824_1.fastq.gz
wget -nc ftp://ftp.sra.ebi.ac.uk/vol1/fastq/ERR377/003/ERR3773823/ERR3773823_1.fastq.gz
wget -nc ftp://ftp.sra.ebi.ac.uk/vol1/fastq/ERR377/008/ERR3773808/ERR3773808_1.fastq.gz
wget -nc ftp://ftp.sra.ebi.ac.uk/vol1/fastq/ERR377/002/ERR3773822/ERR3773822_2.fastq.gz
wget -nc ftp://ftp.sra.ebi.ac.uk/vol1/fastq/ERR377/003/ERR3773823/ERR3773823_2.fastq.gz
wget -nc ftp://ftp.sra.ebi.ac.uk/vol1/fastq/ERR377/009/ERR3773819/ERR3773819_2.fastq.gz
wget -nc ftp://ftp.sra.ebi.ac.uk/vol1/fastq/ERR377/004/ERR3773824/ERR3773824_2.fastq.gz
wget -nc ftp://ftp.sra.ebi.ac.uk/vol1/fastq/ERR377/008/ERR3773818/ERR3773818_2.fastq.gz
wget -nc ftp://ftp.sra.ebi.ac.uk/vol1/fastq/ERR377/006/ERR3773826/ERR3773826_2.fastq.gz
wget -nc ftp://ftp.sra.ebi.ac.uk/vol1/fastq/ERR377/005/ERR3773825/ERR3773825_2.fastq.gz
wget -nc ftp://ftp.sra.ebi.ac.uk/vol1/fastq/ERR377/001/ERR3773811/ERR3773811_1.fastq.gz
wget -nc ftp://ftp.sra.ebi.ac.uk/vol1/fastq/ERR377/003/ERR3773813/ERR3773813_1.fastq.gz
wget -nc ftp://ftp.sra.ebi.ac.uk/vol1/fastq/ERR377/009/ERR3773809/ERR3773809_2.fastq.gz
wget -nc ftp://ftp.sra.ebi.ac.uk/vol1/fastq/ERR377/009/ERR3773819/ERR3773819_1.fastq.gz
wget -nc ftp://ftp.sra.ebi.ac.uk/vol1/fastq/ERR377/007/ERR3773807/ERR3773807_2.fastq.gz
wget -nc ftp://ftp.sra.ebi.ac.uk/vol1/fastq/ERR377/002/ERR3773812/ERR3773812_1.fastq.gz
wget -nc ftp://ftp.sra.ebi.ac.uk/vol1/fastq/ERR377/008/ERR3773808/ERR3773808_2.fastq.gz
wget -nc ftp://ftp.sra.ebi.ac.uk/vol1/fastq/ERR377/000/ERR3773820/ERR3773820_2.fastq.gz
wget -nc ftp://ftp.sra.ebi.ac.uk/vol1/fastq/ERR377/008/ERR3773818/ERR3773818_1.fastq.gz
wget -nc ftp://ftp.sra.ebi.ac.uk/vol1/fastq/ERR377/004/ERR3773814/ERR3773814_1.fastq.gz
wget -nc ftp://ftp.sra.ebi.ac.uk/vol1/fastq/ERR377/001/ERR3773821/ERR3773821_2.fastq.gz
wget -nc ftp://ftp.sra.ebi.ac.uk/vol1/fastq/ERR377/005/ERR3773815/ERR3773815_1.fastq.gz
wget -nc ftp://ftp.sra.ebi.ac.uk/vol1/fastq/ERR377/007/ERR3773817/ERR3773817_1.fastq.gz
wget -nc ftp://ftp.sra.ebi.ac.uk/vol1/fastq/ERR377/006/ERR3773816/ERR3773816_1.fastq.gz
```

```
############################################################
#                        Ragnarok!                         #
############################################################
```

```
#!/bin/bash
#SBATCH -J nf_annot
#SBATCH -A acf-utk0032
#SBATCH --partition=long
#SBATCH --qos=long
#SBATCH --nodes=1
#SBATCH --cpus-per-task=4
```

```
#SBATCH --mem=10G
#SBATCH --time=3-00:00:00
#SBATCH --error=logs/job.e%J
#SBATCH --output=logs/job.o%J
#SBATCH --mail-type=END,FAIL
#SBATCH --mail-user=

source config

cds=$DATA/reference/maize/Zm-B73-REFERENCE-NAM-5.0_Zm00001eb.1.canonical.cds.fa
genome=$DATA/reference/maize/Zm-B73-REFERENCE-NAM-5.0.fa
protein=$DATA/reference/maize/Zm-B73-REFERENCE-NAM-5.0_Zm00001eb.1.protein.fa
ill=$DATA/raw/maize/ERR

nextflow  ~/nextflow/ragnarok/main.nf \
   --design         ~/nextflow/ragnarok/assets/mikado_conf.tsv \
   --publish_dir    $RESULTS/maize_ragnarok \
   --genome         $genome \
   --protein        $protein \
   --ill            $ill \
   --scoring        $DATA/db/plant.yaml \
   --homology       $DATA/db/uniprotkb_taxonomy_id_33090_AND_reviewe_2025_03_07.fasta \
   --skip_qc        false \
   --skip_trim      false \
   --nlrs           false \
   --perform_masking true \
   --cds            $cds \
   --contam         "insecta,fungi,bacteria" \
   -profile         slurm,custom \
   -resume


#############################################################
#                          BRAKER3                          #
#############################################################

#!/bin/bash
#SBATCH -J nf_annot
#SBATCH -A acf-utk0032
#SBATCH --partition=long
#SBATCH --qos=long
#SBATCH --nodes=1
#SBATCH --cpus-per-task=60
#SBATCH --mem=100G
#SBATCH --time=4-00:00:00
#SBATCH --error=logs/job.e%J
#SBATCH --output=logs/job.o%J
#SBATCH --mail-type=END,FAIL
#SBATCH --mail-user=

source config

genome=$DATA/reference/maize/edta_masked_Zm-B73-REFERENCE-NAM-5.0.fa
```

```
protein=$DATA/reference/maize/Zm-B73-REFERENCE-NAM-5.0_Zm00001eb.1.protein.fa

export AUGUSTUS_CONFIG_PATH=$DERIVED/config

singularity exec -B $PROJ docker://teambraker/braker3:v3.0.7.6 \
  braker.pl \
    --threads 48 \
    --workingdir=$RESULTS/maize_braker3 \
    --AUGUSTUS_CONFIG_PATH=$DERIVED/config \
    --genome=$genome \
    --species=maize_new \
    --prot_seq=$protein \
    --
rnaseq_sets_ids=ERR3773807,ERR3773808,ERR3773809,ERR3773810,ERR3773811,ERR3773812,E
RR3773813,ERR3773814,ERR3773815,ERR3773816,ERR3773817,ERR3773818,ERR3773819,ERR377
3820,ERR3773821,ERR3773822,ERR3773823,ERR3773824,ERR3773825,ERR3773826,ERR3773827 \
    --rnaseq_sets_dirs=$DATA/raw/maize/braker_SRA
```

```
############################################################
#                        comparisons                        #
############################################################

#!/bin/bash
#SBATCH -J nf_annot
#SBATCH -A acf-utk0032
#SBATCH --partition=short
#SBATCH --qos=short
#SBATCH --nodes=1
#SBATCH --cpus-per-task=40
#SBATCH --mem=20G
#SBATCH --time=0-03:00:00
#SBATCH --error=logs/job.e%J
#SBATCH --output=logs/job.o%J
#SBATCH --mail-type=END,FAIL
#SBATCH --mail-user=

source config

braker_gtf=$RESULTS/maize_braker3/braker.gtf
b73_gff=$DATA/reference/maize/Zm-B73-REFERENCE-NAM-5.0_Zm00001eb.1.gff3

# convert braker3 gtf to gff3 for use with mikado util stats
singularity exec -B $PROJ docker://quay.io/biocontainers/agat:1.4.2--pl5321hdfd78af_0 \
  agat_convert_sp_gxf2gxf.pl \
    -g $braker_gtf \
    -o $RESULTS/maize_braker3/braker.gff3

# correct the gff for use with mikado, keep all gene models, and all features
singularity exec -B $PROJ docker://quay.io/biocontainers/gffread:0.12.7--h077b44d_6 \
```

```
  gffread \
  -O \
  -E \
  -o $DERIVED/gffread_Zm-B73-REFERENCE-NAM-5.0_Zm00001eb.1.gff3 \
  --keep-genes \
  $b73_gff

#!/bin/bash
#SBATCH -J nf_annot
#SBATCH -A acf-utk0032
#SBATCH --partition=short
#SBATCH --qos=short
#SBATCH --nodes=1
#SBATCH --cpus-per-task=40
#SBATCH --mem=20G
#SBATCH --time=0-03:00:00
#SBATCH --error=logs/job.e%J
#SBATCH --output=logs/job.o%J
#SBATCH --mail-type=END,FAIL
#SBATCH --mail-user=

source config

out_dir=$RESULTS/annotation_metrics
mkdir -p $out_dir

braker_gff=$RESULTS/maize_braker3/braker.gff3
ragnarok_gff=$RESULTS/maize_ragnarok/publish/mikado_final/mikado.loci_out.entap_fil
tered.gff3
b73_gff=$DERIVED/gffread_Zm-B73-REFERENCE-NAM-5.0_Zm00001eb.1.gff3

singularity exec -B $PROJ docker://gemygk/mikado:v2.3.5rc2 \
  mikado util stats $braker_gff $out_dir/braker3_stats.txt

singularity exec -B $PROJ docker://gemygk/mikado:v2.3.5rc2 \
  mikado util stats $ragnarok_gff $out_dir/ragnarok_stats.txt

singularity exec -B $PROJ docker://gemygk/mikado:v2.3.5rc2 \
  mikado util stats $b73_gff $out_dir/b73_stats.txt

#!/bin/bash
#SBATCH -J annot_vs
#SBATCH -A acf-utk0032
#SBATCH --partition=short
#SBATCH --qos=short
#SBATCH --nodes=1
#SBATCH --cpus-per-task=40
#SBATCH --mem=20G
```

```
#SBATCH --time=0-03:00:00
#SBATCH --error=logs/job.e%J
#SBATCH --output=logs/job.o%J
#SBATCH --mail-type=END,FAIL
#SBATCH --mail-user=

source config

out_dir=$RESULTS/annotation_metrics
mkdir -p $out_dir

braker_gff=$RESULTS/maize_braker3/braker.gff3
ragnarok_gff=$RESULTS/maize_ragnarok/publish/mikado_final/mikado.loci_out.entap_fil
tered.gff3
b73_gff=$DERIVED/gffread_Zm-B73-REFERENCE-NAM-5.0_Zm00001eb.1.gff3

singularity exec -B $PROJ docker://gemygk/mikado:v2.3.5rc2 \
   mikado compare \
      -r $b73_gff \
      -p $ragnarok_gff \
      -o $out_dir/compare_ragnarok_maizegdb_stats.txt \
      -x 30

singularity exec -B $PROJ docker://gemygk/mikado:v2.3.5rc2 \
   mikado compare \
      -r $b73_gff \
      -p $braker_gff \
      -o $out_dir/compare_braker3_maizegdb_stats.txt
      -x 30
```
